# Structure-activity relationship of 2,4-D correlates auxin activity with the induction of somatic embryogenesis in *Arabidopsis thaliana*

**DOI:** 10.1101/2022.08.17.504315

**Authors:** Omid Karami, Hanna de Jong, Victor J. Somovilla, Beatriz Villanueva Acosta, Aldo Bryan Sugiarta, Tom Wennekes, Remko Offringa

## Abstract

2,4-dichlorophenoxyacetic acid (2,4-D) is a synthetic analogue of the plant hormone auxin that is commonly used in many *in vitro* plant regeneration systems, such as somatic embryogenesis (SE). Its effectiveness in inducing SE, compared to the natural auxin indole-3-acetic acid (IAA), has been attributed to the stress triggered by this compound rather than its auxin activity. However, this hypothesis has never been thoroughly tested. Here we used a library of 40 2,4-D analogues to test the structure-activity relationship with respect to the capacity to induce SE and auxin activity in *Arabidopsis thaliana*. Four analogues induced SE as effectively as 2,4-D and 13 analogues induced SE but were less effective. Based on root growth inhibition and auxin response reporter expression, the 2,4-D analogues were classified into different groups, ranging from very active auxins to not active. A halogen at the 4-position of the aromatic ring was important for auxin activity, whereas a halogen at the 3-position resulted in reduced activity. Moreover, a small substitution at the carboxylate chain was tolerated, as was extending the carboxylate chain with two but not with one carbon. In the process, we also identified two 2,4-D analogues as efficient inducers of adventitious root formation and several possible anti-auxins. The auxin activity of the 2,4-D analogues was consistent with their simulated TIR1-Aux/IAA coreceptor binding characteristics. A strong correlation was observed between SE induction efficiency and auxin activity, indicating that the stress-related effects triggered by 2,4-D that are considered important for SE induction are down-stream of auxin signaling.

## Introduction

The plant hormone auxin plays a central role in the development of plants. In the 1930s, the structure of the natural auxin indole-3-acetic acid (IAA) was first described (Ma *et al.*, 2018). A few years later, during WWII, 2,4-dichlorophenoxyacetic acid (2,4-D) was discovered as a synthetic auxin analogue that can be used as a herbicide, targeting dicots. Today, it is still broadly used as such in gardens and in agriculture (Peterson *et al.*, 2016). Apart from the cell elongation promoting effect, which led to its discovery as auxin analogue, 2,4-D acts differently in various physiological and molecular assays compared to the natural auxin IAA (Peterson *et al.*, 2016; Calderón Villalobos *et al.*, 2012; Shimizu-Mitao and Kakimoto, 2014).

Binding of IAA to its receptors triggers a transcriptional response. The classical nuclear IAA signaling pathway relies on the degradation of the transcriptional AUXIN/INDOLE-3-ACETIC ACID (Aux/IAA) repressors, leading to expression of auxin responsive genes (Leyser, 2018). The degradation of Aux/IAAs is initiated by binding of auxin to the F-Box proteins TRANSPORT INHIBITOR RESISTANT1 (TIR1) or AUXIN SIGNALING F-BOX 1-3 (AFB1-3). Auxin acts as a molecular glue, allowing the TIR1/AFBs to recruit the N-terminal domain II of Aux/IAAs (Winkler *et al.*, 2017). TIR1/AFBs are part of a Skp1-Cullin-F-box (SCF) E3 ubiquitin ligase complex, which after recruiting Aux/IAAs marks these proteins for degradation by the 26S proteasome (Iglesias *et al.*, 2018; dos Santos Maraschin *et al.*, 2009).

The synthetic auxin 2,4-D elicits a dual response in plants, as it acts as an auxin and also induces stress (Song, 2014; Karami and Saidi, 2010). Like IAA, 2,4-D acts through the TIR1/AFB auxin-mediated signaling pathway (Calderón Villalobos *et al.*, 2012; Shimizu-Mitao and Kakimoto, 2014; Song, 2014). At high concentrations, 2,4-D acts as herbicide and selectively kills dicot weeds. The herbicidal activity of 2,4-D can be attributed to several effects, including the altering of cell wall plasticity and the increase of ethylene levels (Song, 2014). The overproduction of ethylene triggers the increased production of abscisic acid (ABA), which contributes to stomatal closure and thus eventually to plant death (Song, 2014; Karami and Saidi, 2010). 2,4-D also induces the production of reactive oxygen species (ROS) that are very toxic to the plant (Karami and Saidi, 2010).

Besides its use as a herbicide, 2,4-D is widely used for biological experiments to induce auxin responses and for *in vitro* regeneration systems. Our interest in 2,4-D lies in its ability to efficiently induce somatic embryogenesis (SE) at non-herbicidal concentrations. SE is a unique developmental process in which differentiated somatic cells can acquire totipotency and are ‘reprogrammed’ to form new ‘somatic’ embryos. SE is a powerful tool in plant biotechnology used for clonal propagation, genetic transformation and somatic hybridization. It prevents somaclonal variation that is often observed in plant tissue culture, and somatic embryos can be easily cryopreserved (Guan *et al.*, 2016; Horstman *et al.*, 2017). Although SE can be induced by IAA, other synthetic auxins, or stress, in many cases 2,4-D is most effective and it is therefore widely used for SE in many plant species (Karami and Saidi, 2010; Fehér, 2015). Different observations have led to the suggestion that signaling pathways activated by abiotic stress treatments and 2,4 □D treatment converge to regulate a common downstream pathway (Fehér, 2015). 2,4 □D induced SE is accompanied by upregulation of stress related genes (Salvo *et al.*, 2014; Jin *et al.*, 2014; Nowak *et al.*, 2015), and a number of stress related transcription factors have been shown to influence the progression of SE (Gliwicka *et al.*, 2013; Nowak *et al.*, 2015; Mantiri *et al.*, 2008). This suggests that stress induced by non-herbicidal concentrations of 2,4-D (Fehér, 2015) contributes significantly to its effectiveness as SE inducer. To further understand 2,4-D-induced SE, we established a structure-activity relationship (SAR) of 2,4-D with respect to its capacity to induce SE and activity as auxin analogue.

Previous studies have already discovered several chemical analogues of 2,4-D with auxin-like activity. Unfortunately, however, several different bioassays have been used in these studies (Ferro *et al.*, 2010), making a direct comparison difficult. In addition, a number of possible 2,4-D analogues that based on these previous studies would be interesting candidates to assess in a structure-activity relationship (SAR) have not been tested yet. Here we, therefore formed a rationally designed library of 40 2,4-D analogues and screened them for SE induction and auxin activity in *Arabidopsis thaliana* (Arabidopsis). Four analogues induced SE as effectively as 2,4-D and 10 analogues did induce SE but were less effective. Based on root growth inhibition, root hair and lateral root induction, and auxin response reporter expression we classified compounds as very active, active, weakly active or inactive auxins, or even having anti-auxin activity. From the SAR, we concluded that a halogen at the 4-position of the aromatic ring is important for auxin activity, whereas a halogen at the 3-position has a negative effect. Moreover, a small substitution at the carboxylate chain, a methyl or ethyl group, is tolerated, as is extending the carboxylate chain with two but not with one carbon. Molecular dynamics simulations indicated that the classification of the 2,4-D analogues reflected the binding characteristics (enthalpy and distance) to TIR1. SE induction capacity clearly correlated with auxin activity, indicating that the stress response induced by 2,4-D treatment and reported to be required for SE induction is downstream of auxin signaling.

## Results

### Assembly of the 2,4-D analogue library

We aimed to screen a library of 2,4-D analogues for SE induction and auxin activity in *Arabidopsis thaliana* and thereby gain insight into the SAR for this process. To this end, we first performed a literature survey of key past auxin studies in order to identify known 2,4-D analogues for our library (Simon and Petrášek, 2011; Lee *et al.*, 2014; Ferro *et al.*, 2007; Katekar, 1979; Porter and Thimann, 1965; Song, 2014; Tan *et al.*, 2007; van der Zaal *et al.*, 1996; Hayashi *et al.*, 2012; Hamilton *et al.*, 1952; Quareshy *et al.*, 2018; Koepfli *et al.*, 1938; Ferro *et al.*, 2010). This provided an overview of auxin activities and the binding efficiencies of some 2,4-D-like compounds to the auxin co-receptors and gave some context on the SAR of auxin-like compounds. In addition, it offered insight into the difficulties encountered in the past in establishing a SAR, such as non-standardized assays and the use of impure compounds. However, a rational SAR of 2,4-D could not be deduced from these past studies. To achieve a more complete SAR of 2,4-D with respect to the induction of SE and auxin activity, we assembled our own library with TIR1 as the target co-receptor, 2,4-D as a lead compound, analogue hits from the literature survey and several additional structures to achieve a more complete and rational coverage of possible 2,4-D analogues. In general, analogues of the lead compound 2,4-D, either commercially available or synthesized (Supplementary Information), were selected based on modifications at two positions: the type and number of substituents on the aromatic ring or on the alpha position of the carboxylate side chain (**F**ig. 1A). A complete overview of the library is provided in Fig. 1B.

**Figure 1.**
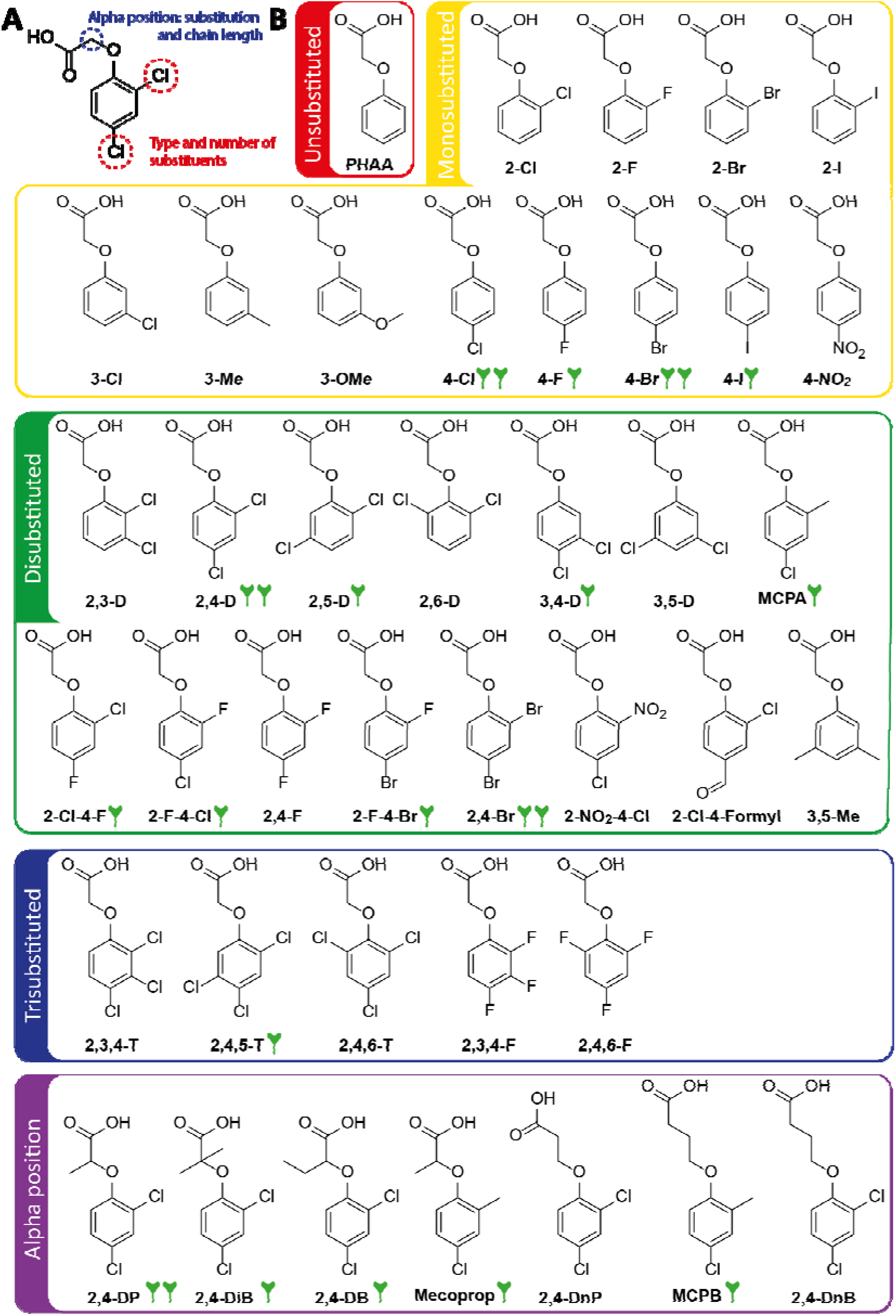
The composition of the 2,4-D analog library. (A) Structure of 2,4-D with the different modifications indicated that compose the 2,4-D analogue library. (B) Structures of the 2,4-D analogs categorized based on the number of groups on the aromatic ring, or the substitution or chain length of the alpha position of the carboxylic acid side chain. Analogs that induced somatic embryogenesis efficiently or at a reduced efficiency are marked with 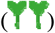 or 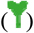, respectively.

### Somatic embryogenesis induction by 2,4-D analogues

As a first step, we tested the ability of all 2,4-D analogues (Fig. 1) to induce SE in *Arabidopsis* immature zygotic embryos (IZEs), which in prior studies has proven to be the most competent *Arabidopsis* tissue for SE in response to 2,4-D (Gaj, 2001).

In this assay, four compounds (class 1: 4-Cl, 4-Br, 2,4-DP, 2,4-Br) were able to efficiently induce somatic embryos from IZEs to a similar extent as 2,4-D (Fig. 2A,B). Thirteen other compounds (class 2: 2,4-DB, 2-Cl-4-F, MCPA, Mecoprop, 4-F, 4-I, 2,5-D, 3,4-D, 2-F-4-Cl, 2-F-4-Br, 2,4,5-T, 2,4-DiB and MCPB) did induce somatic embryos on IZEs but were less effective compared with 2,4-D (Fig. 2A,B). The remaining auxin analogues (class 3: 2-Cl, 2-I, 3-Cl, 2,6-D, 3,5-D, 2,4-F, 2,3,4-F, 2,3,4-T, 2-NO_2_-4-Cl, 2,4-DnB, PHAA, 4-NO_2_, 3,5-Me, 2-Cl-4-formyl, and 2,4,6-F, 3-Me, 2,3-D, 3,5-D, 2,4,6-T and 2,4-DnP) did not induce SE (Fig. S1). Overall, the capacity of compounds to induce SE seemed to correlate with their 2,4-D-like structure.

**Figure 2.**
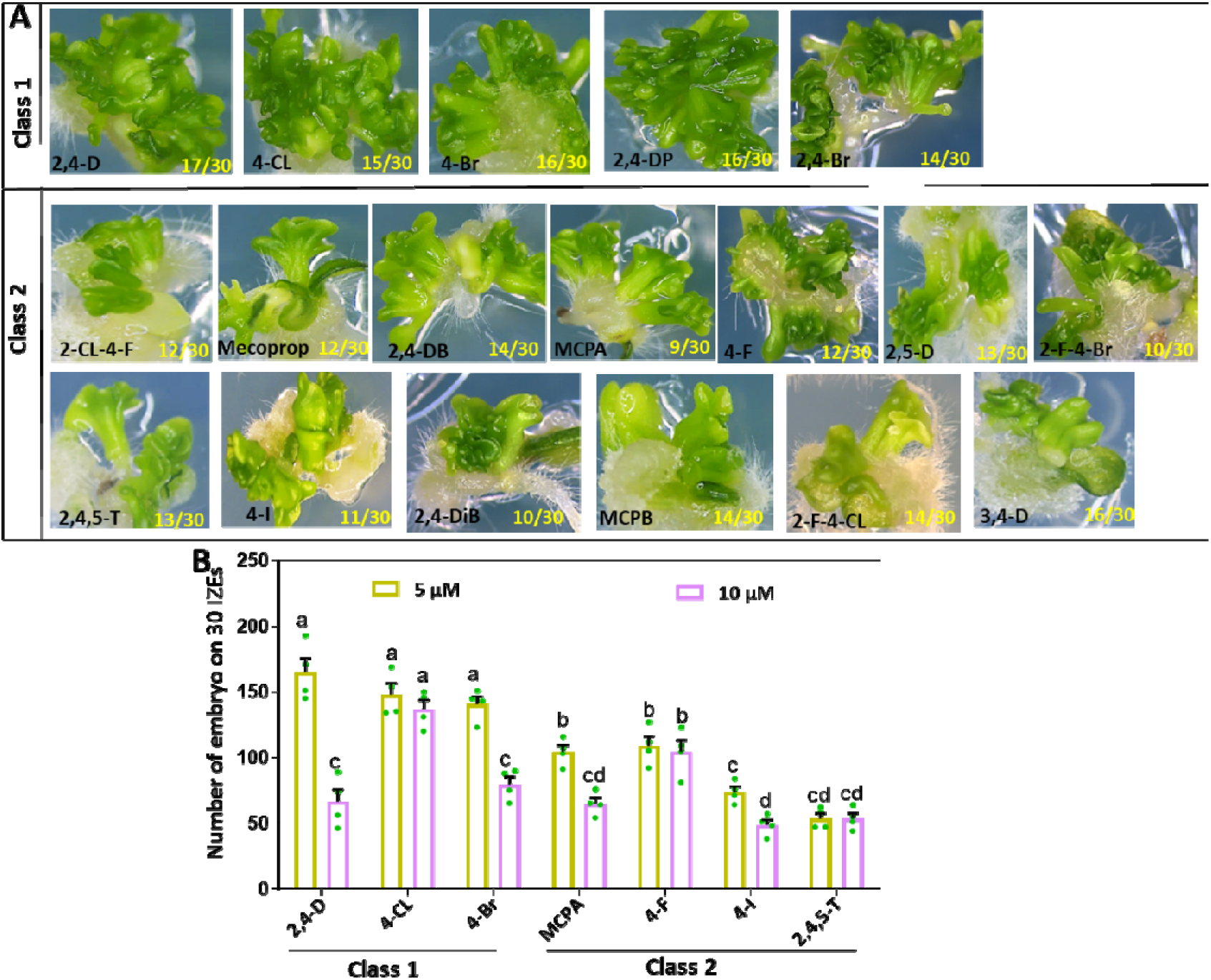
SE inducing capacity of 2,4-D analogues. (A) The phenotype of somatic embryos formed on cotyledons of *Arabidopsis* IZEs that were first grown for two weeks in the presence of 5 μM 2,4-D or one of the 2,4-D analogues and subsequently cultured for 1 week on medium without any supplement. Class 1 compounds induced SE at the same efficiency as 2,4-D at a concentration of 5 μM and class 2 compounds do induce SE but are less effective. (B) Quantification of the number of somatic embryos per 30 IZEs induced by 5 μM or 10 μM of 2,4-D or of the indicated class 1 or class 2 2,4-D analogues. Dots indicate the number somatic embryos produced per IZE (n = 4 biological replicates, with 30 IZEs per replicate), bars indicate the mean value and error bars the SEM. Different letters indicate statistically significant differences (P < 0.001) as determined by one-way analysis of variance with Tukey’s honest significant difference post hoc test. The structure of each compound is provided in **Figure 1**.

We also found that a higher concentration (10 μM) of 2,4-D, 4-Br, 4-I and MCPA reduced the number of embryos formed on IZEs, whereas 10 μM of 4-Cl, 4-F, 2,4,5-T and NAA had no significant effect on the SE efficiency (Fig. 2B). This different SE response to higher levels of certain 2,4-D analogues is probably associated with different metabolic properties, such as uptake and active transport, of these compounds in plant cells.

### Assessing auxin activity of the 2,4-D analogues based on root growth inhibition

Exogenous auxin inhibits primary root elongation in *Arabidopsis* in a concentration-dependent manner (Rahman *et al.*, 2007). As a first assay to establish auxin activity of the 2,4-D analogues in our library (Fig. 1), we tested the inhibitory effect of low (50 nM) and high (5 μM) concentrations on the primary root growth of *Arabidopsis* seedlings (Fig. 3). Similar to 2,4-D, the compounds 4-Cl, 4-Br, 2-Cl-4-F, 2,4-DB, MCPA and Mecoprop inhibited the primary root elongation efficiently, both at low and high concentrations (Fig. 3). Hence, we classified these compounds in one group called very active auxins.

**Figure 3.**
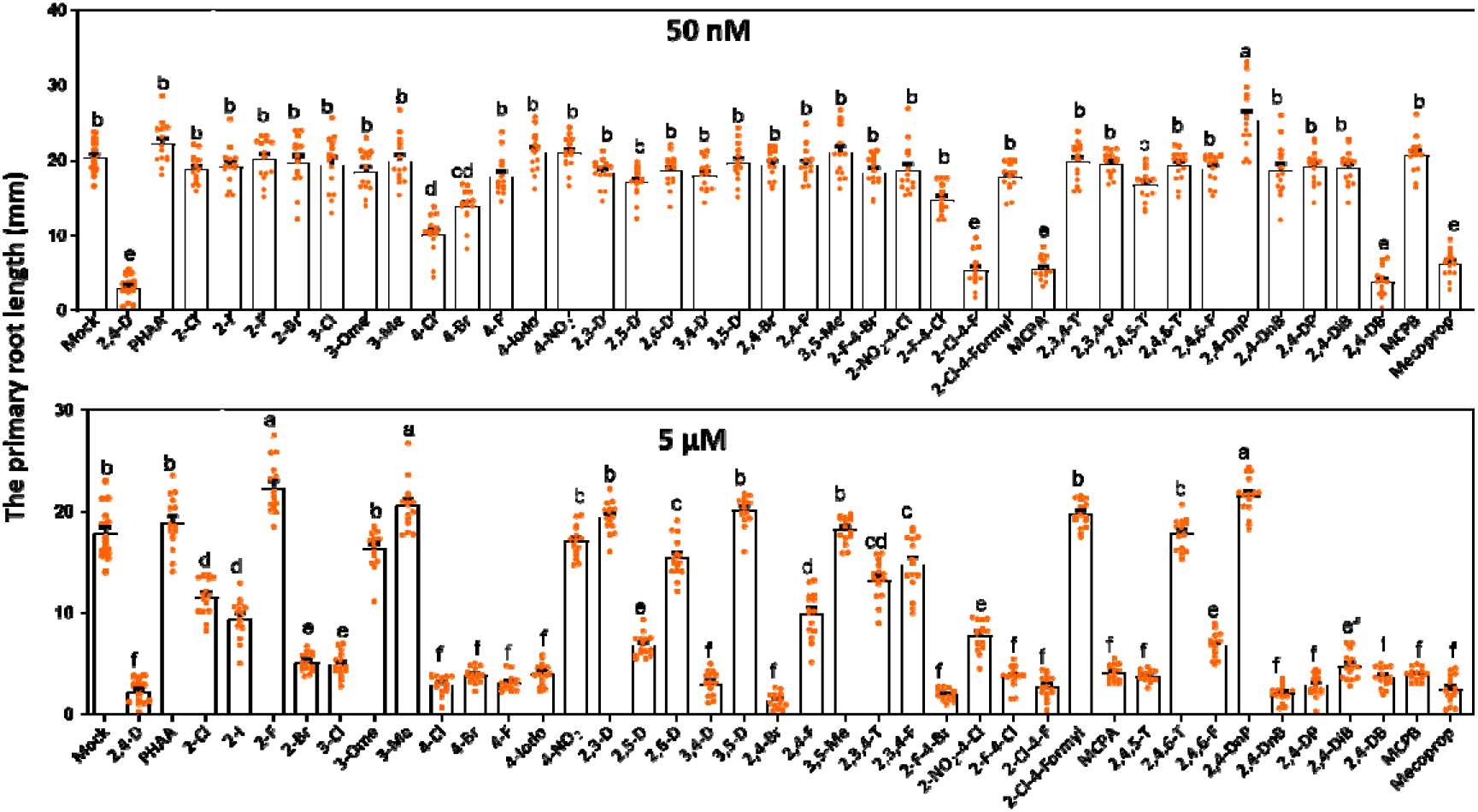
Effect of 2,4-D structure modifications on primary root growth inhibition in *Arabidopsis*. The primary root growth inhibition of seedlings grown in the presence of 50 nM (upper graph) and 5 μM (lower graph) of 2,4-D and 2,4-D analogues. Seedlings were first grown for 6 days on compound free-medium and subsequently transferred on medium with 2,4-D or a 2,4-D analogue for 3 days. Dots indicate the root length (mm) (n=15 biological replicates), bars indicate the mean and error bars the s.e.m. and different letters indicate statistically significant differences (P < 0.01) as determined by a one-way analysis of variance with Tukey’s honest significant difference post hoc test. The structure of each compound is provided in **Figure 1**.

Several compounds (4-F, 4-I, 2,5-D, 3,4-D, 2,4-Br, 2-F-4-Cl, 2-F-4-Br, 2,4,5-T, 2,4-DiB and 2,4-DP, MCPB) showed no inhibitory effect at 50 nM (Fig. 3), whereas they exhibited a strong inhibitory effect at 5 μM (Fig. 3). These compounds were classified as active auxins. By contrast, 2-Cl, 2-I, 3-Cl, 2,6-D, 2,4-F, 2-NO_2_-4-Cl, 2,3,4-T, 2,3,4-F, 2,4,6-T, 2,4,6-F, 2,4-DnP and 2,4-DnB had only a weak effect on root growth at 5 μM (Fig. 3) and were therefore classified as weak auxins. The remaining compounds PHAA, 2-F, 3-Me, 2,3-D, 3,5-M, 3-OMe, 4-NO_2_, 3,5-Me, 3,5-D, 2-CL-4-Formyl, 2,4,6-T, and 2,4-DnP had no obvious inhibitory effect on root length, even at 5 μM (Fig. 3), suggesting that these compounds have either very weak or no auxin activity (Fig. 3). For a complete overview of the 2,4-D analogs categorized based on their auxin activity, the reader is referred to the supplementary data (Fig. S2). The above data indicate that 2,4-D analogues inhibit root growth in both dose- and structure-dependent manner.

### The auxin response induced by 2,4-D analogues corresponds to their root growth inhibition activity

The *pDR5* promoter is a synthetic, generic auxin-responsive promoter that has been extensively used to visualize the cellular auxin response in *Arabidopsis* tissues (Ulmasov *et al.*, 1997). To determine whether the root growth inhibition by 2,4-D analogues correlated with a molecular auxin response, we used an *Arabidopsis* line containing *pDR5* fused to a glucuronidase (GUS) reporter (*pDR5:GUS*). 2,4-D itself induced *pDR5* efficiently, as observed by a completely blue root following histochemical staining for GUS activity. (Fig. 4A). Because 2,4-D induced an especially strong *pDR5* activity in the cell division area of the root tip, we next quantified the effect of the 2,4-D analogues on *pDR5* activity in this part of the root tip (Fig. 4B).

**Figure 4.**
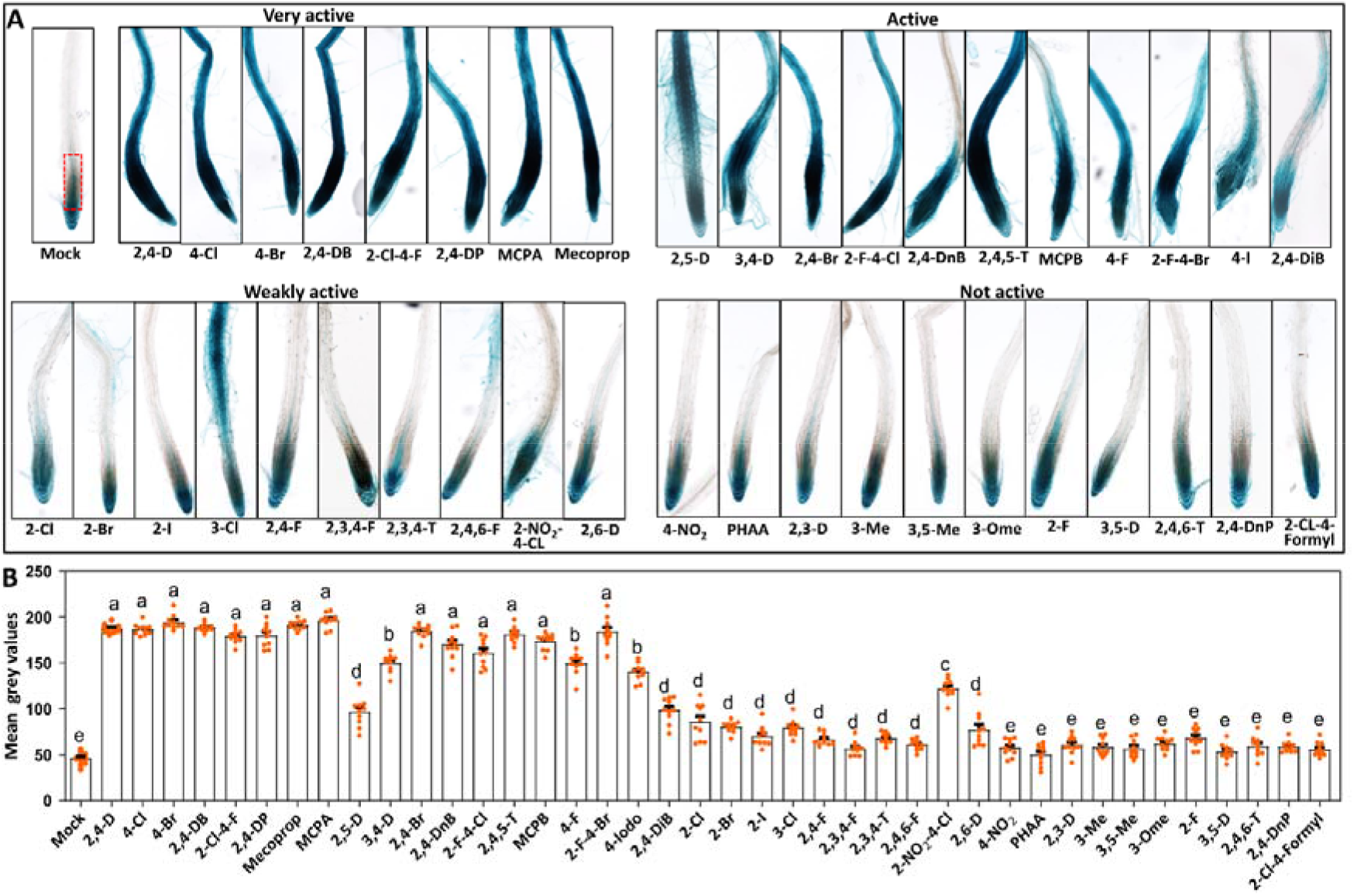
Auxin response induced by 2,4-D analogues. (A) Expression of the auxin responsive *pDR5:GUS* reporter in *Arabidopsis* seedling roots treated for 24 hours with 5 μM of each 2,4-D analogue. For an overview of the 2,4-D analogues and their abbreviations, see **Fig. 2**. (B) Quantification of *pDR5:GUS* activity in the root meristems of *Arabidopsis* seedlings (region indicated with dotted red line in mock). Dots indicate the individual measurement of GUS activity staining per each treatment (n= 10 biologically independent seedlings per treatment, bars indicate the mean of each treatment and error bars the s.e.m. and different letters indicate statistically significant differences (P < 0.01) as determined by a one-way analysis of variance with Tukey’s honest significant difference post hoc test. In A and B, seedlings were germinated for 6 days on compound-free medium, and for treatment seedlings were transferred to medium contain 5 μM of a 2,4-D analogue and incubated for 24 hours. The structure of each compound is provided in **Figure 1**.

As expected, the compounds classified as very active auxins (Fig. S2) all strongly promoted *pDR5* activity in the cell elongation zone of *Arabidopsis* roots (Fig. 4A, B), in line with their root growth inhibition activity. The *pDR5:GUS* expression was also clearly induced but varied more for the active auxins (Fig. 4A, B). The compounds that had only a slight effect on root growth and were therefore classified as weak auxins, also only slightly promoted expression of the *pDR5:GUS* reporter (Fig. 4A, B). Interestingly, 3-Cl induced a stronger auxin response along the differentiation zone of the root (Fig. 4A) compared to the other compounds classified as weak auxins. The remaining library members that did not lead to inhibition of root growth, also did not promote *pDR5* activity (Fig. 4A, B). These findings support our conclusion that the ability of the 2,4-D analogues to inhibit root growth correlates well with their auxin response induction.

To further confirm the auxin activity of the 2,4-D analogues, we used the nuclear auxin input reporter R2D2, which acts as a proxy for the cellular sensing of auxin (Liao *et al.*, 2015). The R2D2 reporter consists of two parts, an auxin-degradable DII domain fused to the yellow fluorescent nVENUS that is rapidly degraded when auxin concentrations increase as auxin sensor and an auxin-nondegradable DII domain (mDII) fused to the orange fluorescent nTdTOMATO as expression control (Liao *et al.*, 2015). In accordance with observations based on *pDR5* activity, we detected strong down-regulation of the DII-nVENUS signal in the cell elongation zone of *Arabidopsis* roots treated with all very active and most of the active 2,4-D analogues (2,5-D, 3,4-D, 2,4-Br, 2-F-4-Br, 2,4-DP, and MCPB) and two weakly active compounds (2-Br and 2-NO_2_-4-Cl) (Fig. S3). The other active and weakly active compounds elicited only moderate downregulation of DII-Venus (Fig. S3). The compounds that proved inactive as an auxin, as these did not inhibit root growth nor promoted *pDR5* activity, also did not lead to a detectable reduction of the DII-nVENUS signal in the cell elongation zone of roots (Fig. S3). These results again supported the overall classification of the auxin activity for our library of 2,4-D analogues (Fig. S2).

### Specific 2,4-D analogues as tools to modulate root system architecture

When establishing the root growth inhibition of our 2,4-D analogues, we observed that several compounds had a unique effect on the root system architecture (RSA). The RSA of a plant describes the organization of the primary, lateral and adventitious roots. This includes root hairs that increase the surface area and thus promote the uptake of water and nutrients (Smith and de Smet, 2012). Auxin treatment is known to change the RSA by inducing lateral or adventitious roots (Hayashi *et al.*, 2008; da Costa *et al.*, 2018) and to positively influence the formation of root hairs in a dose-dependent manner (Lee and Cho, 2013). An adventitious root (AR) refers to a plant root that forms from any non-root tissue, commonly in response to treatment with the natural auxin indole-3-butyric acid (IBA) (Geiss *et al.*, 2009; da Costa *et al.*, 2018).

All 2,4-D analogues that were designated as very active, active and weakly active positively increased the number of root hairs in Arabidopsis seedling roots (not shown). Specifically, we observed a strong effect on root hair formation on root tissues treated with 5 μM of 3-Cl, 3,4-D, 2,4-DP or Mecoprop. We found that the root hair formation was dose-dependent for all four analogues (Fig. S4). Interestingly, treatment with 0,5 or 1 μM of the weakly active 3-Cl did not lead to a strong inhibition of root growth, like with the other three active or very active compounds (Fig. S4), whereas it still strongly enhanced root hair development. This result reflects the weaker auxin response induced by 3-Cl in the root meristem, leading to reduced root growth inhibition (Fig. 2), while it still induces a relatively strong auxin response in the differentiation zone, resulting in ectopic root hair formation (Fig. 4).

We also observed that 3-Cl and 2,5-D significantly promoted the number of lateral roots, whereas 2,4-D and other analogues with strong auxin activity initially induced many lateral root meristems, but these meristems quickly deteriorated into amorphous callus (Fig. 5A). As lateral root and adventitious root (AR) induction are highly linked, we tested the capacity of different concentrations of 2,4-D, IBA, 3-Cl and 2,5-D in AR induction from hypocotyls of dark grown *Arabidopsis* seedlings. AR induction is a crucial process in clonal crop propagation by cuttings or shoot regeneration, and is well-known to be promoted in many plant species by the natural auxin indole-3-butyric acid (IBA). As with lateral roots and in line with previous observations (da Costa *et al.*, 2018), treatment with 2,4-D produced a low number of ARs at low μM concentrations and only undesired callus at higher μM concentrations (Fig. 5B,C). Treatment with 3-Cl and 2,5-D, however, efficiently induced ARs at 5 μM (Fig. 5B, C). In addition, the number of ARs induced by 3-Cl and 2,5-D was significantly higher compared to IBA treatment at a similar concentration and also compared to 57 μM IAA or 2 μM of the synthetic auxin 1-naphthaleneacetic acid (NAA) (Fig. 5D), treatments that have previously been shown to efficiently induces ARs from of *Arabidopsis* hypocotyls (da Costa *et al.*, 2018). With these results, we can conclude that 3-Cl and 2,5-D are excellent candidates for inducing AR formation in Arabidopsis, and that despite their structural similarity with 2,4-D, they show a unique biological activity. This probably relates to their mild and specific activity as auxin analogue, as reflected by the specific expression pattern of the *pDR5:GUS* reporter following treatment with these compounds.

**Figure 5.**
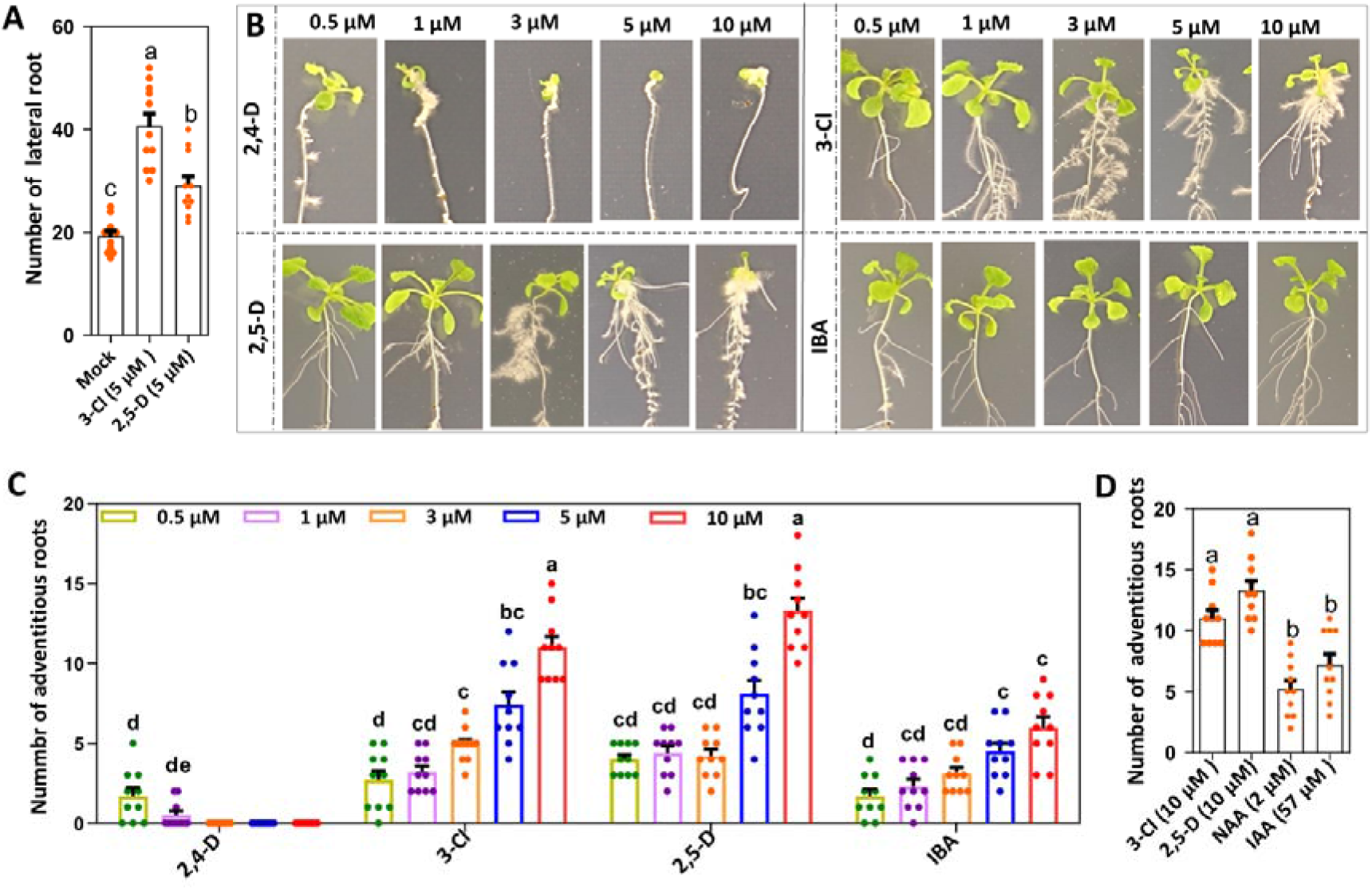
Efficient adventitious root induction on etiolated *Arabidopsis* hypocotyls by specific 2,4-D analogues. (A) Number of lateral roots formed in Arabidopsis seedlings treated with mock or 5 μM of 3-Cl and 2,5-D. Seedlings were germinated for 6 days on compound-free medium, and subsequently transferred for treatment to medium containing 5 μM of a 2,4-D analogue and incubated for 10 days. (B) The phenotype adventitious roots induced on hypocotyls of 6-day-old etiolated Arabidopsis seedlings by transferring them to medium with different concentrations of 2,4-D, 2,5-D, 3-Cl or IBA. (C) Quantification of the number of adventitious roots induced on etiolated *Arabidopsis* hypocotyls by 2,4-D, 2,5-D, 3-Cl or IBA (see B). (D) Comparison of the number of adventitious roots induced from etiolated *Arabidopsis* hypocotyls by 10 μM of 2,5-D, 10 μM of 3-Cl, 2 μM of NAA or 57 μM of IAA. In A, C and D dots indicate the number lateral roots per main root in A and the number adventitious roots per hypocotyl in C and D (A. n=15; C. n=10; D. n=10), the bars indicate the mean value, and the error bars the s.e.m.. The letters above the bars indicate the significant difference (p < 0.01) determined by a one-way ANOVA with Tukey’s honest significant difference post hoc test. In B-D, *Arabidopsis* seedlings were initially cultured for 6 days on hormone-free medium in the dark, and subsequently the seedlings were transferred on medium containing the indicated compound.

Lateral root formation can be repressed by anti-auxins (Hayashi *et al.*, 2008; Larsen, 2017). Several of the 2,4-D analogues for which we observed no clear effect on root length or *pDR5* activity, namely 3-Me, 2,3-D, 3,5-D, 2,4,6-T, and 2,4-DnP, did significantly inhibit lateral root formation, with some having a stronger effect (Fig. 6A, B). In order to investigate the lateral root inhibition-responsiveness to these potential anti-auxins, we examined the effect of short term treatment (1 day) on the different developmental stages of lateral root formation, using the *pDR5:GUS* reporter activity as marker (Dubrovsky *et al.*, 2008). Histochemical analysis revealed that *pDR5:GUS* is expressed in the presence of anti-auxins in all lateral root stages, the lateral root initiation (stage 1), lateral root development (stage 2), lateral root emergence (stage 3) and lateral root outgrowth (stage 4) (not shown). This is in line with the observation that treatment with these compounds did not lead to a reduced expression of the *pDR5:GUS* reporter in the main root tip (Fig. S2). However, these 2,4-D derived anti-auxin candidates did reduce the number of stage 1 primordia, had no effect on the number of stage 3 primordia, whereas they had a differential effect on the number of stage 2 primordia and stage 4 lateral roots (Fig. 6C). The effect of these compounds on lateral root formation might indeed be caused by their activity as anti-auxins, however we cannot exclude that they affect other processes, such as auxin transport.

**Figure 6.**
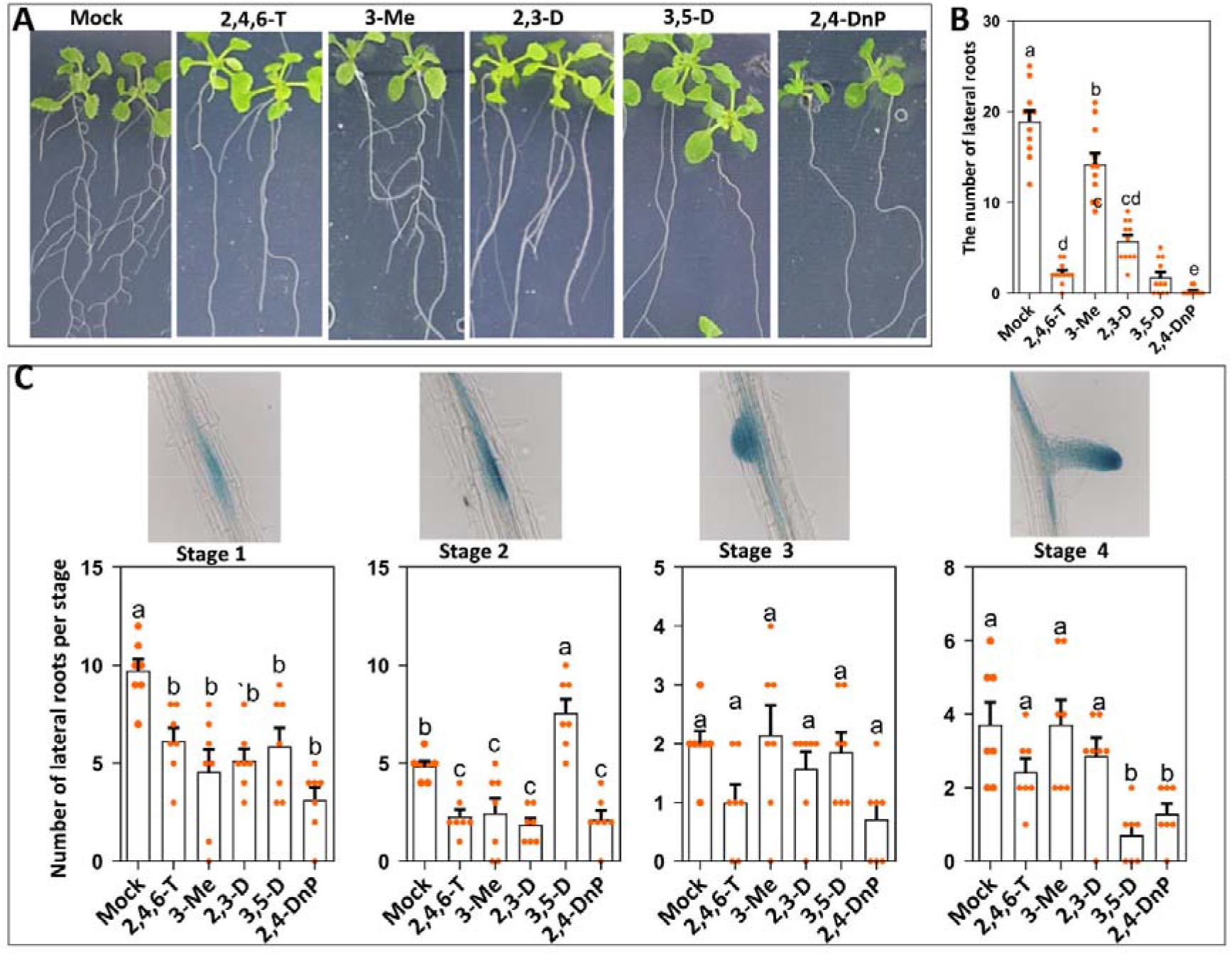
Several 2,4-D analogues inhibit lateral root formation in *Arabidopsis*. (A) Root phenotype of 16-day-old Arabidopsis seedlings grown on medium containing the indicated 2,4-D analogue. (B) The number of clearly visible lateral roots formed on *Arabidopsis* seedling primary roots treated with mock or 5 μM of the indicated 2,4-D analogue. (C) Quantification of the effect of 1 day treatment on the different developmental stages of lateral root formation including lateral root initiation (stage 1), lateral root primordium development (stage 2), lateral root emergence (stage 3) and lateral root outgrowth (stage 4 and longer) using *pDR5:GUS* activity to visualize the early stages of lateral root development. In B and C dots indicate the number of lateral roots in B and number of lateral roots per developmental stage in C (B: n=10 and C: n=7), the bars indicate the mean value and the error bars the standard error. The letters above the bars indicate the significant difference (p < 0.01) determined by a one-way ANOVA with Tukey’s honest significant difference post hoc test. Seedlings were first grown for 6 days on hormone-free medium, and subsequently transferred to mock medium or medium with 5 μM of the indicated compound and further incubated for 10 days in A and B and for 1 day in C.

Based on the combined results we now classified all tested 2,4-D analogues from our library into five groups, namely very active, active, weakly active auxin analogues and not active and anti-auxin compounds (Fig. S2). Our results show that the capacity of these auxin analogues to induce SE is tightly linked to their auxin activity.

### Molecular dynamics simulations of binding of 2,4-D analogues to the TIR1-Aux/IAA coreceptors

To correlate the auxin activity of the 2,4-D analogues to their binding to the TIR1/AFB – Aux/IAA auxin co-receptors (Tan *et al.*, 2007), we performed molecular dynamics (MD) simulations of this binding for a subset of 14 2,4-D analogues, representing derivatives from all auxin activity classes as determined in the *in planta* experiments. 2,4-D, IAA and NAA were taken along as controls (Table 1). Among the library entries tested *in silico* are alpha substituted compounds (methyl, ethyl or the Mecoprop derivatives 2,4-DP, 2,4-DB and Mecoprop, respectively) that due to their stereocenter can occur as the enantiomeric R or S stereoisomers. These compounds were tested as mixture of these enantiomers in the *in planta* assays. The molecular dynamics simulation uniquely allowed us to investigate at a molecular level the potential differential binding of each enantiomer (notated as their acronym with an added R and S).

**Table 1.** Activity of auxins and 2,4-D analogues compared to the parameters from the molecular dynamics simulations for their binding to the TIR1-Aux/IAA coreceptors. The auxins and 2,4-D analogs are ordered according to their established auxin activity of the SAR (see also Table S2). Several parameters were calculated from the molecular dynamics simulations: i) the enthalpy binding of the selected compounds and the standard deviation (std) was determined, ii) the root-mean-square deviation of the TIR1 protein (Protein RMSD) with a standard deviation (std) was calculated as a measure of system stability during the simulation trajectory, iii) the distance between the gamma carbon of the proline of the AUX/IAA peptide and the ring mass center of the compound and the corresponding standard deviation were determined (Distance gamma). At short distances, a stronger CH·π interaction is assumed. A larger distances (5Å and more) the CH·π contribution is more or less negligible (No CH π)

| Compound | Auxin activity | Enthalpy binding<br>(Kcal/mol) | std | Protein RMSD (Å) | std | Distance gamma<br>(Å) | std |
| --- | --- | --- | --- | --- | --- | --- | --- |
| 2,4-D | very active | -26,9 | 0,84 | 1,4 | 0,12 | 4,2 | 0,38 |
| IAA | very active | -26,8 | 0,80 | 1,2 | 0,13 | 4,0 | 0,28 |
| NAA | very active | -29,6 | 1,06 | 1,6 | 0,15 | 4,3 | 0,39 |
| 2,4-DB(S) | very active | -30,8 | 0,70 | 1,3 | 0,16 | 4,3 | 0,33 |
| 2,4-DB(R) | very active | -29,3 | 0,86 | 1,3 | 0,12 | 4,5 | 0,45 |
| Mecoprop(R) | very active | -26,7 | 0,70 | 1,5 | 0,19 | 3,7 | 0,3 |
| Mecoprop(S) | very active | -15,1 | 0,70 | 1,3 | 0,1 | 4,3 | 0,36 |
| 2,4-DP(R) | very active | -25,1 | 1,51 | 1,5 | 0,11 | 4,0 | 0,37 |
| 2,4-DP(S) | very active | -22,4 | 0,65 | 1,2 | 0,1 | 4,2 | 0,33 |
| 2-Cl-4-F | very active | -23,2 | 0,81 | 1,3 | 0,12 | 4,0 | 0,31 |
| 4-Cl | very active | -13,0 | 0,53 | 1,4 | 0,15 | 3,9 | 0,28 |
| MCPA | very active | -12,8 | 0,69 | 1,4 | 0,15 | 3,8 | 0,29 |
| 2,4-DiB | active | -19,5 | 0,43 | 1,5 | 0,14 | No CH·π | 0,53 |
| 3,4-D | active | -7,8 | 0,54 | 1,3 | 0,09 | 4,0 | 0,37 |
| 2,6-D | weakly active | -36,5 | 0,76 | 1,4 | 0,13 | No CH·π | 0,43 |
| 2,3,4-T | weakly active | -8,8 | 0,57 | 1,2 | 0,11 | 3,6 | 0,23 |
| 2,3,4-F | weakly active | -8,3 | 0,67 | 1,3 | 0,11 | 4,2 | 0,47 |
| 4-NO<sub>2</sub> | not active | -13,6 | 0,54 | 1,5 | 0,23 | No CH·π | 0,96 |
| 2,4-DnP | anti-auxin | -8,4 | 1,06 | 1,1 | 0,09 | 4,2 | 0,57 |
| 2,3-D | anti-auxin | -6,6 | 0,69 | 1,3 | 0,13 | 4,2 | 0,55 |

In the evaluation of the binding capacity of each auxin analogue within the system, we studied different aspects: the enthalpy of the system, the root-mean-square deviation (RMSD) of atomic positions of TIR1 along the simulation trajectory, the hydrogen bond network established by the auxin analogue and the distance between the auxin analogue and the proline (7) of the Aux/IAA peptide (Table 1). The RMSD of TIR1 reflects the system stability during the simulation. The last parameter, the distance (gamma) between the auxins analogue and the proline (7) of the Aux/IAA peptide, would show us which analogue is able to establish a CH·π interaction with the degron peptide that contributes to the binding interactions and thus to the stability of the complex (Wang and Yao, 2019). Our molecular dynamics simulations were based on a previously published method (Hao and Yang, 2010), but with a much longer trajectory of 200 ns instead of 2.5 ns.

The enthalpy was calculated using MMPBSA (Molecular Mechanics Poisson-Boltzmann Surface Area) with a lower enthalpy value indicating better binding. All the compounds classified as very active auxin analogues showed an enthalpy value below −12 Kcal/mol. Surprisingly, the weakly active 2,6-D and inactive 4-NO_2_ were also in this range, suggesting that the binding enthalpy is not the sole parameter predictive for auxin activity. The TIR1 protein RMSD calculation was carried out over the amino acids from position 50 till the end of the protein, because the first 50 amino acids showed extra flexibility. The TIR1 N-terminal part is normally stabilized by the interaction with the SKP protein, which we did not include in our analysis. Moreover this part does not interact with the auxin analogue nor with the AUX/IAA peptide and its structure has not been completely resolved in the employed crystal structure (Tan *et al.*, 2007). All the compounds induced similar RMSD values of the TIR1 protein ranging from 1.25 to 1.53, indicating that binding to the compounds tested do not induce significant differences in the stability of the protein backbone (Table 1). All compounds tested appeared to establish a hydrogen bond network but with different effectiveness (Fig. S5). Serine (438) and arginine (403) interact through a hydrogen bond with all the derivatives, but these interactions have different prevalence. Other amino acids that are involved in most of the hydrogen bond networks established by the analogues are arginine (436) and histidine (78). Finally, we observed the proximity between the proline (7) from the AUX/IAA peptide and the analogue, so we monitored this distance along the simulation trajectory for all the derivatives. The distance was measured specifically between the gamma carbon of proline (7) and the ring mass center of the auxin analogue (distance gamma), assuming that a shorter distance would predict a stronger CH·π interaction (Wang and Yao, 2019). Compounds 4-NO_2_ (5,3 Å) and 2,6-D (5,0 Å) showed the two largest distances at which the CH·π contribution is more or less negligible, explaining why these compounds are weakly active or inactive despite their low enthalpy. For IAA and the very active 2,4-D analogues distance values varied between 3,7 Å (Mecoprop) and 4,5 Å (2,4-DB(R)). Several weakly active and anti-auxin compounds were also in this range. However, as the binding enthalpy was above −12 Kcal/mol, this would explain why these compounds lacked auxin activity.

Summarizing, there is a strong correlation between auxin activity and SE induction: SE is only obtained with the very active and active auxin analogs and the very active auxins are generally more efficient in inducing SE than the active auxins (Fig. S2).

## Discussion

Chemical biology has been instrumental in enhancing our understanding of auxin biology, as several key molecules that are currently used to specifically control auxin biosynthesis, signaling or transport have been identified in chemical genetic screens (Simon *et al.*, 2013; Ma *et al.*, 2018). 2,4-D is a synthetic auxin that is widely used as a herbicide, but also as a plant growth regulator in *in vitro* regeneration and auxin research (Skùpa *et al.*, 2014; Peterson *et al.*, 2016). In this study, 2,4-D analogues with varying modifications, including the type and number of substitutions on the aromatic ring (like chlorine, bromine, fluorine, iodine, nitro, methyl group) or substitutions on or elongation of the alpha-position (Fig. 1) have been evaluated for their structure-activity relationship and ability to induce SE.

### The capacity of SE induction by 2,4-D analogues

SE is experimentally induced by application of exogenous plant growth regulators under *in vitro* conditions (Nic-Can *et al.*, 2015). 2,4-D is the most well-known plant growth regulator that is widely used for SE. Our screen revealed four 2,4-D analogues (4-Cl, 4-Br, 2,4-DP, and 2,4-Br) that also efficiently induce SE from Arabidopsis IZEs, whereas 13 analogues (4-F, 4-I, 2,5-D, 3,4-D, 2-F-4-Cl, 2-F-4-Br, 2,4,5-T, 2,4-DiB, MCPB, MCPA, and Mecoprop) induced SE at a lower efficiency (Fig. 2 and Fig. S1). Except for 3,4-D and 2,4,5-T, which have been shown to efficiently induce SE in other plant species (Kouassi *et al.*, 2017), SE induction by the other 2,4-D analogues has not been reported before. We did not identify a 2,4-D analogue with enhanced capacity to induce SE, suggesting that for our system 2,4-D is still the best compound to use. However, since 2,4,5-T efficiently induces SE in other plant species but less efficiently in Arabidopsis, it is to be expected that some of the 2,4-D analogues presented here might be used to establish an efficient SE system in other plant species. Moreover, 2,4-D treatment has been reported to lead to a significant percentage of abnormal somatic embryos in many plant species (Garcia *et al.*, 2019). The use of the 2,4-D analogues identified here may reduce the number of abnormal embryos, but this still requires testing in the different SE systems.

We also showed that only 2,4-D analogues that classify as very active or active auxins are able to induce SE from Arabidopsis IZEs, which indicates that the capacity to induce SE is primarily linked to auxin activity. Since stress has also been identified as an important factor in SE induction (Mantiri *et al.*, 2008; Salvo *et al.*, 2014; Jin *et al.*, 2014; Nowak *et al.*, 2015), our findings suggest that at least in our SE system stress is downstream of auxin signaling, and not a parallel pathway that is additionally triggered by 2,4-D or its analogues. It is therefore unlikely that a chemical biology approach as presented here will allow to tease apart the stress and auxin pathways by identifying compounds that trigger each pathway separately. The new 2,4-D analogues identified here can be useful tools to study the importance of aspects of auxin physiology either during SE induction or through their effects on the root system architecture. For example, it would be interesting to understand why MCPA, which is classified as very active auxin based on root growth inhibition and auxin response reporter expression, shows a significantly reduced capacity to induce SE. And why does 3-Cl, classified as weakly active auxin, strongly induce the auxin response in the root differentiation zone thereby specifically promoting root hair formation? Or how can 3,5-D strongly inhibit lateral root formation without clearly affecting auxin responsive gene expression? These differences may lie in the compound specific metabolism or transport characteristics of the 2,4-D analogues, which has already been shown to differ between the natural auxin IAA and the synthetic auxins 2,4-D and 1-NAA (Delbarre *et al.*, 1996; Song, 2014; Eyer *et al.*, 2016).

### Auxin activity of 2,4-D analogues correlates to simulated binding properties to TIR1

The Arabidopsis seedling root growth inhibition assay combined with the use of the *pDR5:GUS* and *R2D5* auxin response reporters provided consistent results with respect to classifying the 40 2,4-D orthologs as very active, active or weakly active auxins, or having no auxin activity. Importantly, this classification was in agreement with previously published results on the 2,4-D analogues PAA, 2,4 Br, 2,4,5-T, 3-Me, 2,4-DP and MCPB (Simon *et al.*, 2013; Torii *et al.*, 2018). This difference in auxin activity is clearly determined by structure and chemical properties of these compounds, which relates on the one hand to their binding affinity to the TIR1/AFB (Calderón Villalobos *et al.*, 2012) and AUX/IAA (Torii *et al.*, 2018) co-receptor pair, and on the other hand to their transport or conjugation properties or metabolic decay in plant cells. For example, 2,4-D itself it has been shown to be conjugated to Asp and Glu, and the conjugates still display residual auxin activity (Eyer *et al.*, 2016).

Molecular dynamics simulations indicated that the auxin activity of 2,4-D analogues could be correlated to their binding strength to TIR1. One important factor here was the theoretical enthalpy of the system, as all very active auxin analogues showed an enthalpy value below −12 Kcal/mol. However, the distance between Pro7 of Aux/IAA and the 2,4-D analogue was also important, as some of the tested analogues lacking this CH·π interaction showed no or only weak auxin activity, despite the fact that their enthalpy value was below −12 Kcal/mol.

It is important to note that, due to the expensive cost of entropy calculation combined with low reliability for such a big system, we did not calculate this parameter. However, we considered that the molecules are very similar and, although we know there will be differences in entropy among the derivatives, we assume that they will not change the results dramatically.

### Structure-activity relationship of 2,4-D analogues based on auxin activity

Based on our screen of 40 compounds, we have identified several trends for the structure auxin activity relationship of 2,4-D analogues. First, a halogen at the 4-position is more important for auxin activity than at the 2-position (4-Cl; 2-Cl-4-F; 4-I/Br/F; versus 2-Cl/I/Br/F). Second, the activity of analogues with a substitution at the 3- or 5-position remains, but such compounds are less active compared to 2,4-substitutions (2,5-D; 3,4-D; 2,4,5-T). Third, both substitutions at the alpha position (Mecoprop; 2,4-DP; 2,4-DB; 2,4-DiB) and a longer carboxylate chain with an even number of carbons are tolerated (MCPB; 2,4-DnB). This finding is in agreement with a previous study that showed that IAA analogues with a substitution on their carboxylate chain, up to n=4 carbons, can still bind to TIR1, thus allowing for a modification on this position (Hayashi *et al.*, 2008). Taken these trends together, we conclude that an electron withdrawing group, in particular a halogen, at the 4-position is important for auxin activity. In addition, we suggest that electron withdrawing or donating properties of the 2-position are less important for auxin activity and that the size of the 2-substituent could influence auxin activity, since both methyl and chloro groups are accepted at this position (like 2,4-D and MCPA). There are, however, a few outliers based on this general conclusion: 2,4-F is only weakly active, but has a strong electron withdrawing group at the 4-position, and 2,4-DnB is also weakly active, but has the same substitution as 2,4-D and a longer carboxylate chain which does not disrupt auxin activity for MCPB. Altogether, these results indicate that small modifications at different parts of 2,4-D can lead to different physiological activities, generating molecules with interesting other applications in plant (tissue) culture than SE, as described below.

### Several 2,4-D analogues are useful tools to modulate the root system architecture

In the assessment of 2,4-D analogues, we have found that some 2,4-D analogues (2,3-D, 3,5-D, 3-Me, 2,4,6-T and 2,4-DnP) either had no effect on or only slightly affected root growth (Fig. 3), whereas they inhibited the formation of lateral roots (Fig. 6). Since inhibition of lateral root formation is typical for compounds with anti-auxin activity (Larsen, 2017), we suspected that they may act as anti-auxins. Such anti-auxin activity could be caused by high affinity binding to only one of the co-receptors, thereby preventing the formation of the F-box-Aux/IAA complex. As such, high concentrations of an anti-auxin would compete effectively with IAA for establishing the TIR1-Aux/IAA complex required for ubiquitination-mediated degradation of the Aux/IAA repressors. Interestingly, our 2,4-D analogues did not change *pDR5* activity, nor did they lead to enhanced or reduced degradation of DII-VENUS in the root tip, which would be expected for an anti-auxin. Moreover, the negative effect on lateral root formation varied per compound. For example, 3-Me only mildly affected lateral root formation, whereas 3,5-D and 2,4-DnP almost completely inhibited this process, with 3,5-D preferably blocking at stage 2 (enhanced number of stage 2 primordia) whereas 2,4DnP resulted in a reduction of all stages. Clearly, further research on these 2,4-D analogues, such as a forward genetic screen for mutants that develop lateral roots when grown on the compound, is required to unravel the exact molecular mechanism by which they repress lateral root formation. As such, these 2,4-D analogues could uncover new components involved in lateral root formation. At the same time, they could be used to control lateral root formation, e.g. to prevent branching during early seedling development to obtain a deeper root system.

Adventitious root (AR) formation is an organogenesis process by which new roots are produced from non-root tissues. AR induction is an important but often rate limiting step in the vegetative propagation of many horticultural and forestry plant species. Generally, AR induction is promoted by auxins, and IBA and NAA are the most common auxins used for this purpose in commercial operations (Geiss *et al.*, 2009). Our results show that 2,5-D and 3-Cl efficiently induce AR from *Arabidopsis* hypocotyls even more efficiently than IBA or NAA. Therefore, 2,5-D and 3-Cl are recommended as new AR inducers and they may be used to resolve rooting in recalcitrant species.

Roots hairs extend from root epidermal cells in the differentiation zone of the root and support plants in nutrient absorption, anchorage, and microbe interactions (Lee and Cho, 2013). Exogenous auxin treatments generally promotes root hair induction and development (Lee and Cho, 2013). We showed that some 2,4-D analogues (3-Cl, 3,4-D, 2,4-DP, and Mecoprop) induced ectopic root hair formation on Arabidopsis roots, while promoting lateral root formation and having a relatively mild effect on root growth compared to 2,4-D itself. Specifically, roots of plants grown on the analogue 3-Cl produced several times more root hairs compared to other analogues. Unlike other 2,4-D analogues, 3-Cl specifically induced a strong auxin response in the differentiation zone of the root and a weaker response in the root elongation zone (Fig. S4), this could explain the differential effect of 3-Cl on root biomass. We recommend 3-Cl as a promising compound for use in horticulture or agriculture to enhance root hair formation and thereby improve crop performance by enhanced ion and water uptake.

## Materials and methods

### 2,4-D and 2,4-D-analogues

The 2,4-D analogues that were used in this study were divided into 2 main categories. The first one contained 2,4-D analogues with substituents at different positions on the aromatic ring (Fig. S1). The second group consisted of 2,4-D analogues with different carbon chains at the alpha-position of 2,4-D. (Fig. S2) For more information about the synthesis of certain library members and the sources for commercially available compounds, the reader is kindly referred to the Supplementary Information.

The acronyms of the 2,4-D analogues were assigned based on established acronyms in literature, or based on the number and substitution type on the aromatic ring, or the type of substituent on the alpha position of the carboxylate.

### Plant materials and growth conditions

This research used *Arabidopsis thaliana* Columbia-0 (Col-0) wild-type, and the *pDR5::GUS* and *R2D2* reporter lines in the Columbia background. For *in vitro* plant culture, seeds were sterilized in 10 % (v/v) sodium hypochlorite for 10 minutes and then 4 times washed with MQ water. Sterilized seeds were plated on ½ MS medium and grown in the tissue culture room at 21°C, 16 hours photoperiod and 50 % relative humidity. After around 2 weeks, the germinated seeds were planted in soil in the climate room at 20°C, 16 hours photoperiod and 70 % relative humidity.

### *Arabidopsis* primary root growth assay adventitious root induction

The *Arabidopsis* seeds were germinated on ½ MS medium. Five-day-old seedlings were transferred to new ½ MS medium supplemented with 2,4-D analogues. The length of the primary root was quantified after 3 days, incubation with 2,4-D analogues. Primary root length and lateral root numbers were analyzed with ImageJ software.

For the adventitious root (AR) induction, the seed was first grown in complete darkness. Six-day-old seedlings were transferred to new ½ MS medium supplemented with 2,4-D analogues 16 hours photoperiod. The number of ARs induced from hypocotyls was quantified after 7 days.

### Histological GUS staining assay

In order to test auxin activity of 2,4-D analogues, the activity *pDR5:GUS* reporter was investigated in the presence of 5 μM 2,4-D analogues. Histochemical β-glucuronidase (GUS) staining was performed as described previously (Anandalakshmi *et al.*, 1998) with some modifications. Samples were submerged in 1-2mL staining solution and incubated for 4 hours at 37°C followed by rehydration through a graded ethanol series 75% - 50% - 25% for 10 minutes each, with 5 minutes incubation between each step. *Arabidopsis* Col-0 on MS medium without any supplement was used as a control.

The tissue-specific GUS staining intensity was quantified as mean grey values by analyzing images of independent samples capturing the same region of interest (ROI) using ImageJ, as previously described by Béziat et al. (Béziat *et al.*, 2017).

### Somatic embryogenesis induction

For SE induction, 11 days old siliques of *Arabidopsis* Columbia-0 wild-type were sterilized in 10 % (v/v) sodium hypochlorite for 10 minutes and then washed with MQ water for 4 times. Sterilized siliques were dissected to acquire IZEs. IZEs were cultured on B5 medium mixed with 2,4-D analogues for 14 days in the culture room at 21°C, 16 hours photoperiod and 50 % relative humidity. The IZEs were then transferred to ½ MS medium for 7 days and the number of SEs was counted under a light microscope. The quality of the obtained SEs was then further examined by moving embryos to the MS media in square petri dishes. Some 2,4-D analogues inactive in inducing SE were combined with IAA or NAA. In this experiment, B5 medium supplemented with 2,4-D used as a control.

### Microscopy

GUS stained tissues were observed and photographed using a Nikon’s Eclipse E800 microscope. Cellular and subcellular localization of DII-VENUS and TdTOMATO proteins was visualized by confocal laser scanning microscope (ZEISS-003-18533) using a 534 nm laser combined with a 488 nm LP excitation and 500-525 nm BP emission filters for DII-VENUS signals or 633 laser and 532 nm LP excitation and 580-600 nm BP emission filters for TdTOMATO signals.

### Molecular dynamics simulations

Molecular dynamics simulations were run using the crystal structure of the systems comprising the TIR1, auxin responsive protein fragment, co-factor inositol hexakisphosphate, and the auxin. In particular, the structure deposited in the PDB under the ID 2P1N (Tan et al., 2007) was used as a starting structure. The auxin derivative was replaced by the different auxin candidates before running the MD simulations. Both the system preparation and the simulations were performed in the AMBER 18 suite software (Case, 2018). An updated version of the protocol described in the Hao and Yang’s article was used (Hao and Yang, 2010). Firstly, the system is neutralized by adding sodium ions and later immersed in a cubic box of 10 Å length in each direction using TIP3P (Mol. Phys, 1988, 94, 803-808) water parameters. The force fields used to obtain topography and coordinates files were ff14SB (Maier et al., 2015) and GAFF (Wang et al., 2004). The first step to start MD simulations is a minimization only of the positions of solvent molecules keeping the solute atom positions restrained and, the second stage minimizes of all the atoms in the simulation cell. Heating the system is the third step raising gradually the temperature 0 to 300 K under a constant volume and periodic boundary conditions. In addition, Harmonic restraints of 10 kcal·mol^-1^ were applied to the solute, and the Berendsen temperature coupling scheme (Berendsen *et al.*, 1984) was used to control and equalize the temperature. The time step was kept at 2 fs during the heating phase. Long-range electrostatic effects were modelled using the particle-mesh-Ewald method (Darden *et al.*, 1993). The Lennard-Jones interactions cut-off was set at 8 A□. An equilibration step for 2 ns with a 2 fs time step at a constant pressureand temperature of 300 K was performed prior the production stage. The trajectory production stage kept the equilibration step conditions and was prolonged for 200 ns longer at 1 fs time step. Besides, the auxin analogs required a previous preparation step where the parameters and charges were generated by using the antechamber module of AMBER, using GAFF force field and AM1-BCC method for charges (Jakalian *et al.*, 2002). In addition, 100 frames evenly separated during the last 5000 steps of the simulation were employed for the enthalpy calculation by using MM-PBSA approximation.

## Supplementary data

The following supplementary data are available online.

Fig S1. Weakly or not active 2,4-D analogues do not induce SE.

Fig S2. An overview of the 2,4-D analogues classified very active, active or weakly active auxins, or not active, or anti-auxins.

Fig S3. R2D2-reported auxin activity of the 40 2,4-D analogues.

Fig S4. Certain 2,4-D analogues specifically promote root hair formation.

Fig S5. The hydrogen bond network between the very active 2,4-D analogues and TIR1.

A list of commercially available 2,4-D analogues.

Chemical synthesis of 2,4-D analogues.

Copies of NMR spectra.

## Acknowledgements

This publication is part of the project StressAuxEmb (with project number 737.016.013) of the research programme Building Blocks of Life, which was (partly) financed by the Dutch Research Council (NWO). Victor J Somovilla is thankful for the financial support to EU Commission (Marie Skłodowska-Curie 840663 to V.J.S.) and Maria de Maeztu Units of Excellence Programme – Grant No. MDM-2017-0720 Ministry of Science, Innovation and Universities.

## Author contributions

O.K., H.J. and V.J.S.: Methodology, Formal analysis and Validation. O.K., H.J., V.J.S., B.V.A. and A.B.S.: Investigation. O.K., H.J., T.W. and R.O.: Conceptualization and Writing – Original Draft. All authors: Writing – Review & Editing. T.W. and R.O.: Supervision and Funding Acquisition.

## Data availability

All data supporting the findings of this study are available within the paper and within the supplementary data published online. Raw data are available from the corresponding author, (Remko Offringa), upon request. Novel materials used and described in the paper are available upon request for non-commercial research purposes.

## References

Anandalakshmi, R., Pruss, G.J., Ge, X., Marathe, R., Mallory, A.C., Smith, T.H. and Vance, V.B. (1998) A viral suppressor of gene silencing in plants. Proc. Natl. Acad. Sci. U. S. A., 95, 13079–13084.

Berendsen, H.J.C., Postma, J.P.M., Gunsteren, W.F. van, DiNola, A. and Haak, J.R. (1984) Molecular dynamics with coupling to an external bath. J. Chem. Phys., 81, 3684–3690.

Béziat, C., Kleine-Vehn, J. and Feraru, E. (2017) Histochemical Staining of β-Glucuronidase and Its Spatial Quantification. In J. Kleine-Vehn and M. Sauer, eds. Plant Hormones: Methods and Protocols. New York, NY: Springer New York, pp. 73–80.

Calderón Villalobos, L.I.A., Lee, S., Oliveira, C. De, et al. (2012) A combinatorial TIR1/AFB–Aux/IAA co-receptor system for differential sensing of auxin. Nat. Chem. Biol., 8, 477–485.

Case, D.A. (2018) Amber 18. Univ. California, San Fr.

Costa, C.T. da, Gaeta, M.L., Araujo Mariath, J.E. de, Offringa, R. and Fett-Neto, A.G. (2018) Comparative adventitious root development in pre-etiolated and flooded *Arabidopsis* hypocotyls exposed to different auxins. Plant Physiol. Biochem., 127, 161–168.

Darden, T., York, D. and Pedersen, L. (1993) Particle mesh Ewald: An N ·log(N) method for Ewald sums in large systems. J. Chem. Phys., 98, 10089–10092.

Delbarre, A., Muller, P., Imhoff, V. and Guern, J. (1996) Comparison of mechanisms controlling uptake and accumulation of 2,4-dichlorophenoxy acetic acid, naphthalene-1-acetic acid, and indole-3-acetic acid in suspension-cultured tobacco cells. Planta, 198, 532–541.

Dubrovsky, J.G., Sauer, M., Napsucialy-Mendivil, S., Ivanchenko, M.G., Friml, J., Shishkova, S., Celenza, J. and Benková, E. (2008) Auxin acts as a local morphogenetic trigger to specify lateral root founder cells. Proc. Natl. Acad. Sci. U. S. A., 105, 8790–8794.

Eyer, L., Vain, T., Pařízková, B., et al. (2016) 2,4-D and IAA amino acid conjugates show distinct metabolism in Arabidopsis. PLoS One, 11, e0159269.

Fehér, A. (2015) Somatic embryogenesis - stress-induced remodeling of plant cell fate. Biochim. Biophys. Acta - Gene Regul. Mech., 1849, 385–402.

Ferro, N., Bredow, T., Jacobsen, H.J. and Reinard, T. (2010) Route to novel auxin: Auxin chemical space toward biological correlation carriers. Chem. Rev., 110, 4690–4708.

Ferro, N., Bultinck, P., Gallegos, A., Jacobsen, H.J., Carbo-Dorca, R. and Reinard, T. (2007) Unrevealed structural requirements for auxin-like molecules by theoretical and experimental evidences. Phytochemistry, 68, 237–250.

Gaj, M.D. (2001) Direct somatic embryogenesis as a rapid and efficient system for in vitro regeneration of Arabidopsis thaliana. Plant Cell. Tissue Organ Cult., 64, 39–46.

Garcia, C., Furtado de Almeida, A.A., Costa, M., Britto, D., Valle, R., Royaert, S. and Marelli, J.P. (2019) Abnormalities in somatic embryogenesis caused by 2,4-D: an overview. Plant Cell. Tissue Organ Cult., 137, 193–212.

Geiss, G., Gutierrez, L. and Bellini, C. (2009) Adventitious Root Formation: New Insights and Perspectives. In Root Development. Oxford, UK: Wiley-Blackwell, pp. 127–156.

Gliwicka, M., Nowak, K., Balazadeh, S., Mueller-Roeber, B. and Gaj, M.D. (2013) Extensive Modulation of the Transcription Factor Transcriptome during Somatic Embryogenesis in Arabidopsis thaliana. PLoS One, 8, e69261.

Guan, Y., Li, S.G., Fan, X.F. and Su, Z.H. (2016) Application of somatic embryogenesis in woody plants. Front. Plant Sci., 7, 938.

Hamilton, R.H., Kivilaan, A. and McManus, J.M. (1952) Biological Activity of Tetrazole Analogues of Indole-3-Acetic Acid and 2,4-Dichlorophenoxyacetic Acid. PLANT Physiol., 35, 136–140.

Hao, G.F. and Yang, G.F. (2010) The role of Phe82 and Phe351 in auxin-induced substrate perception by TIR1 ubiquitin ligase: A novel insight from molecular dynamics simulations. PLoS One, 5, e10742.

Hayashi, K.-I., Tan, X., Zheng, N., Hatate, T., Kimura, Y., Kepinski, S. and Nozaki, H. (2008) Small-molecule agonists and antagonists of F-box protein-substrate interactions in auxin perception and signaling. Proc. Natl. Acad. Sci. U. S. A., 105, 5632–7.

Hayashi, K.I., Neve, J., Hirose, M., Kuboki, A., Shimada, Y., Kepinski, S. and Nozaki, H. (2012) Rational design of an auxin antagonist of the SCFTIR1 auxin receptor complex. ACS Chem. Biol., 7, 590–598.

Horstman, A., Bemer, M. and Boutilier, K. (2017) A transcriptional view on somatic embryogenesis. Regeneration, 4, 201–216.

Iglesias, M.J., Terrile, M.C., Correa-Aragunde, N., et al. (2018) Regulation of SCFTIR1/AFBs E3 ligase assembly by S-nitrosylation of Arabidopsis SKP1-like1 impacts on auxin signaling. Redox Biol., 18, 200–210.

Jakalian, A., Jack, D.B. and Bayly, C.I. (2002) Fast, efficient generation of high-quality atomic charges. AM1-BCC model: II. Parameterization and validation. J. Comput. Chem., 23, 1623–1641.

Jin, F., Hu, L., Yuan, D., Xu, J., Gao, W., He, L., Yang, X. and Zhang, X. (2014) Comparative transcriptome analysis between somatic embryos (SEs) and zygotic embryos in cotton: Evidence for stress response functions in SE development. Plant Biotechnol. J., 12, 161–173.

Karami, O. and Saidi, A. (2010) The molecular basis for stress-induced acquisition of somatic embryogenesis. Mol. Biol. Rep., 37, 2493–2507.

Katekar, G.F. (1979) Auxins: On the nature of the receptor site and molecular requirements for auxin activity. Phytochemistry, 18, 223–233.

Koepfli, J.B., Thimann, K. V. and Went, F.W. (1938) Phytohormones; structure and physiological activity. J. Biol. Chem., 122, 736–780.

Kouassi, M.K., Kahia, J., Kouame, C.N., Tahi, M.G. and Koffi, E.K. (2017) Comparing the effect of plant growth regulators on callus and somatic embryogenesis induction in four elite *Theobroma cacao* L. genotypes. HortScience, 52, 142–145.

Larsen, P.B. (2017) Anti-auxin compounds. U.S. Patent 0073308A1.

Lee, R.D.W. and Cho, H.T. (2013) Auxin, the organizer of the hormonal/environmental signals for root hair growth. Front. Plant Sci., 4, 448.

Lee, S., Sundaram, S., Armitage, L., Evans, J.P., Hawkes, T., Kepinski, S., Ferro, N. and Napier, R.M. (2014) Defining Binding Efficiency and Specificity of Auxins for SCF TIR1/AFB - Aux/IAA Co-receptor Complex Formation. ACS Chem. Biol., 9, 673–682.

Leyser, O. (2018) Auxin signaling. Plant Physiol., 176, 465–479.

Liao, C.Y., Smet, W., Brunoud, G., Yoshida, S., Vernoux, T. and Weijers, D. (2015) Reporters for sensitive and quantitative measurement of auxin response. Nat. Methods, 12, 207–210.

Ma, Q., Grones, P. and Robert, S. (2018) Auxin signaling: a big question to be addressed by small molecules. J. Exp. Bot., 69, 313–328.

Maier, J.A., Martinez, C., Kasavajhala, K., Wickstrom, L., Hauser, K.E. and Simmerling, C. (2015) ff14SB: Improving the Accuracy of Protein Side Chain and Backbone Parameters from ff99SB. J. Chem. Theory Comput., 11, 3696–3713.

Mantiri, F.R., Kurdyukov, S., Lohar, D.P., Sharopova, N., Saeed, N.A., Wang, X.D., Vandenbosch, K.A. and Rose, R.J. (2008) The transcription factor MtSERF1 of the ERF subfamily identified by transcriptional profiling is required for somatic embryogenesis induced by auxin plus cytokinin in Medicago truncatula. Plant Physiol., 146, 1622–1636.

Nic-Can, G.I., Galaz-Ávalos, R.M., De-la-Peña, C., Alcazar-Magaña, A., Wrobel, K. and Loyola-Vargas, V.M. (2015) Somatic embryogenesis: Identified factors that lead to embryogenic repression. A case of species of the same genus. PLoS One, 10, e0126414.

Nowak, K., Wójcikowska, B. and Gaj, M.D. (2015) *ERF022* impacts the induction of somatic embryogenesis in Arabidopsis through the ethylene-related pathway. Planta, 241, 967–985.

Peterson, M.A., McMaster, S.A., Riechers, D.E., Skelton, J. and Stahlman, P.W. (2016) 2,4-D Past, Present, and Future: A Review. Weed Technol., 30, 303–345.

Porter, W.L. and Thimann, K. V. (1965) Molecular requirements for auxin action-I. Halogenated indoles and indoleacetic acid. Phytochemistry, 4, 229–243.

Quareshy, M., Prusinska, J., Kieffer, M., et al. (2018) The Tetrazole Analogue of the Auxin Indole-3-acetic Acid Binds Preferentially to TIR1 and Not AFB5. ACS Chem. Biol., 13, 2585–2594.

Rahman, A., Bannigan, A., Sulaman, W., Pechter, P., Blancaflor, E.B. and Baskin, T.I. (2007) Auxin, actin and growth of the *Arabidopsis thaliana* primary root. Plant J., 50, 514–528.

Salvo, S.A.G.D., Hirsch, C.N., Buell, C.R., Kaeppler, S.M. and Kaeppler, H.F. (2014) Whole transcriptome profiling of maize during early somatic embryogenesis reveals altered expression of stress factors and embryogenesis-related genes. PLoS One, 9, e111407.

Santos Maraschin, F. dos, Memelink, J. and Offringa, R. (2009) Auxin-induced, SCFTIR1-mediated poly-ubiquitination marks AUX/IAA proteins for degradation. Plant J., 59, 100–109.

Shimizu-Mitao, Y. and Kakimoto, T. (2014) Auxin sensitivities of all Arabidopsis Aux/IAAs for degradation in the presence of every TIR1/AFB. Plant Cell Physiol., 55, 1450–1459.

Simon, S., Kubeš, M., Baster, P., Robert, S., Dobrev, P.I., Friml, J., Petrášek, J. and Zažímalová, E. (2013) Defining the selectivity of processes along the auxin response chain: A study using auxin analogues. New Phytol., 200, 1034–1048.

Simon, S. and Petrášek, J. (2011) Why plants need more than one type of auxin. Plant Sci., 180, 454–460.

Skûpa, P., Opatrný, Z. and Petrášek, J. (2014) Auxin Biology: Applications and the Mechanisms Behind. In P. Nick and Z. Opatrny, eds. Applied Plant Cell Biology: Cellular Tools and Approaches for Plant Biotechnology. Berlin, Heidelberg: Springer Berlin Heidelberg, pp. 69–102.

Smith, S. and Smet, I. de (2012) Root system architecture: Insights from *Arabidopsis* and cereal crops. Philos. Trans. R. Soc. B Biol. Sci., 367, 1441–1452.

Song, Y. (2014) Insight into the mode of action of 2,4-dichlorophenoxyacetic acid (2,4-D) as an herbicide. J. Integr. Plant Biol., 56, 106–113.

Tan, X., Calderon-Villalobos, L.I.A., Sharon, M., Zheng, C., Robinson, C. V., Estelle, M. and Zheng, N. (2007) Mechanism of auxin perception by the TIR1 ubiquitin ligase. Nature, 446, 640–645.

Torii, K.U., Hagihara, S., Uchida, N. and Takahashi, K. (2018) Harnessing synthetic chemistry to probe and hijack auxin signaling. New Phytol., 220, 417–424.

Ulmasov, T., Murfett, J., Hagen, G. and Guilfoyle, T.J. (1997) Aux/IAA proteins repress expression of reporter genes containing natural and highly active synthetic auxin response elements. Plant Cell, 9, 1963–1971.

Wang, J., Wolf, R.M., Caldwell, J.W., Kollman, P.A. and Case, D.A. (2004) Development and testing of a general amber force field. J. Comput. Chem., 25, 1157–1174.

Wang, J. and Yao, L. (2019) Dissecting C–H⋯π and N–H⋯π Interactions in Two Proteins Using a Combined Experimental and Computational Approach. Sci. Rep., 9, 20149.

Winkler, M., Niemeyer, M., Hellmuth, A., et al. (2017) Variation in auxin sensing guides AUX/IAA transcriptional repressor ubiquitylation and destruction. Nat. Commun., 8, 15706.

Zaal, B.J. van der, Droog, F.N., Pieterse, F.J. and Hooykaas, P.J. (1996) Auxin-sensitive elements from promoters of tobacco *GST* genes and a consensus as-1-like element differ only in relative strength. Plant Physiol., 110, 79–88.

