## Supplementary material for "Structure-activity relationship of 2,4-D correlates auxin activity with the induction of somatic embryogenesis in *Arabidopsis thaliana*": - (from another version of this preprint: version 1, 2022-08-18)

Supporting information

Table of Contents

Figure S1	S2

Figure S2	S3

Figure S3	S4

Figure S4	S5

Commercially available 2,4-D analogues	S6

Chemical synthesis	S6

Copies of NMR spectra	S12

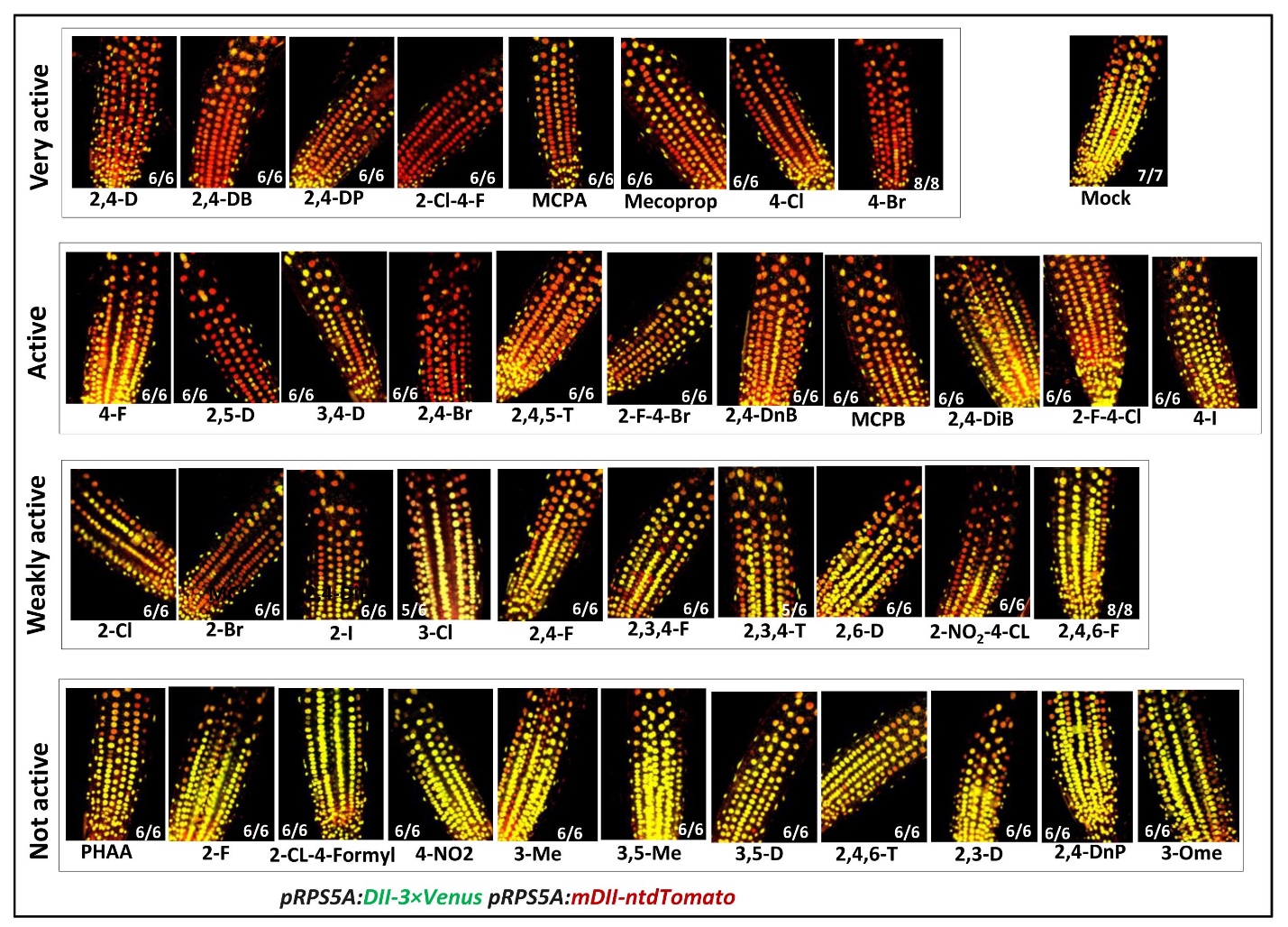

**Figure S1. *R2D2*-reported auxin activity of the 40 2,4-D analogues.** Visualization of the auxin activity of different 2,4-D analogues by using the *DII-Venus/mDII-tomato* (*R2D2*) reporter in the main root meristem of *Arabidopsis thaliana* seedlings treated with the different 2,4-D analogues. A yellow signal is indicative for no or low auxin activity, whereas an orange or red signal indicates intermediate or high auxin activity, respectively. Seeds were first grown for 6 days on control (mock) medium, and subsequently the seeds were transferred to liquid medium containing 5 μM of the 2,4-D analogues for 1 hour.

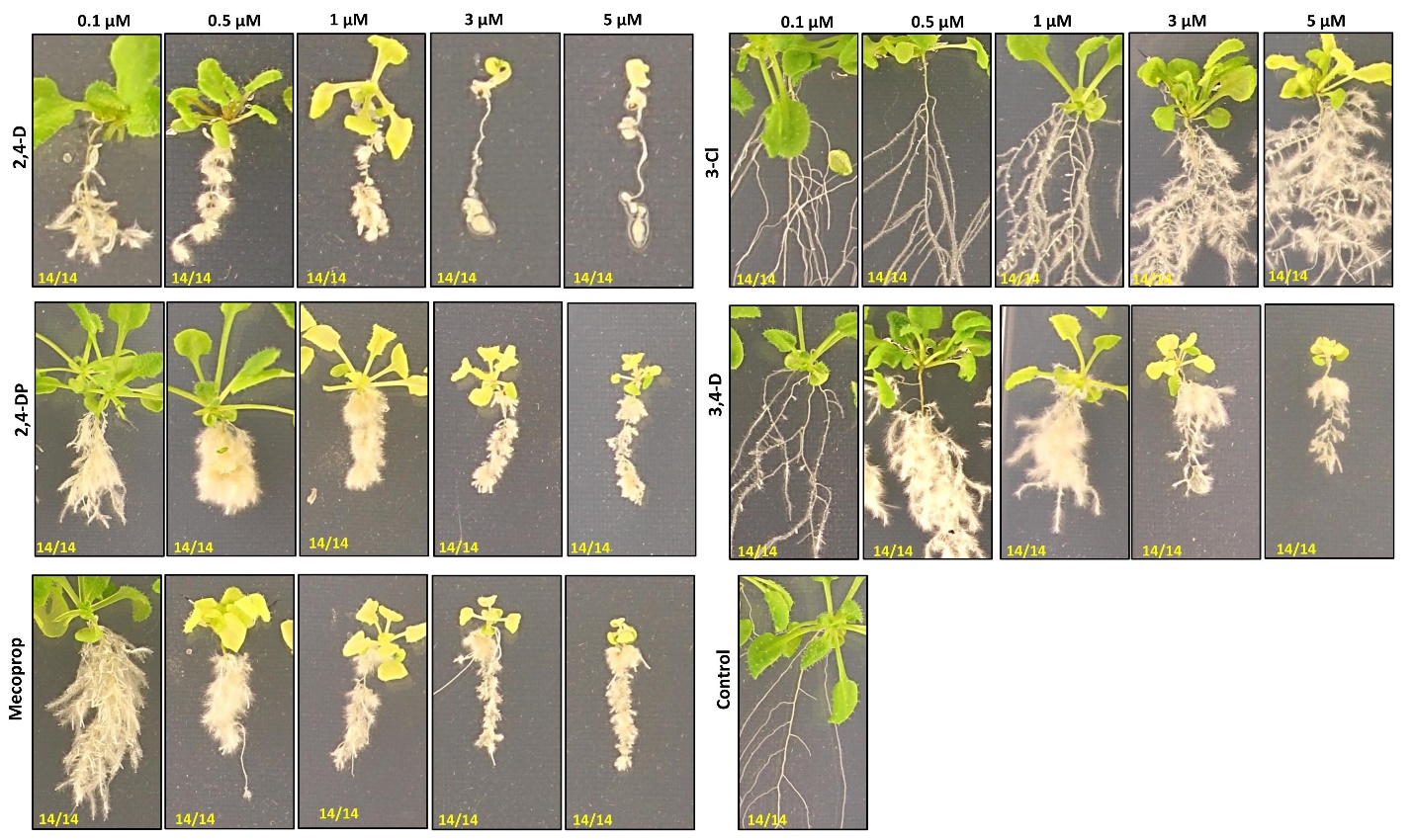

**Figure S2. Certain 2,4-D analogues specifically promote root hair formation.** The phenotype of Arabidopsis seedlings treated with the indicated concentration of 2,4-D, 2,4-DP, Mecoprop, 3-Cl, or 3,4-D. Seedlings were initially grown for 6 days on control medium and subsequently cultured for 15 days on medium containing the indicated concentration of the 2,4-D analogue.

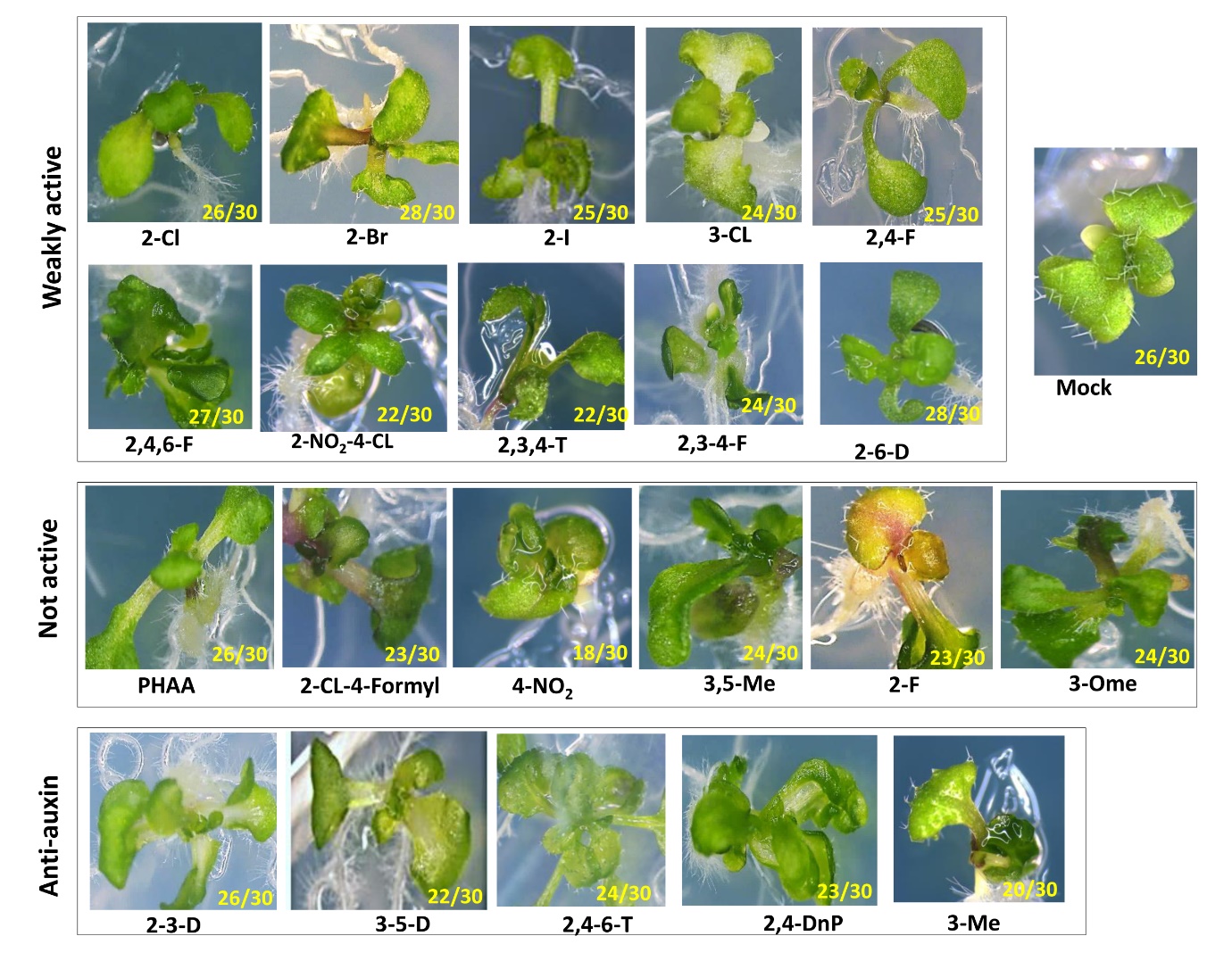

**Figure S3. Weakly or not active 2,4-D analogues do not induce SE.** The phenotype of *Arabidopsis* IZEs that were first grown for two weeks in the presence of 5 μM of the indicated 2,4-D analogue and subsequently cultured for 1 week on medium without any supplement.

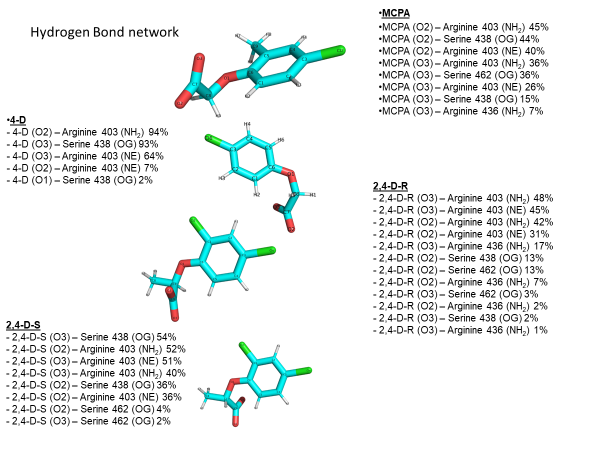

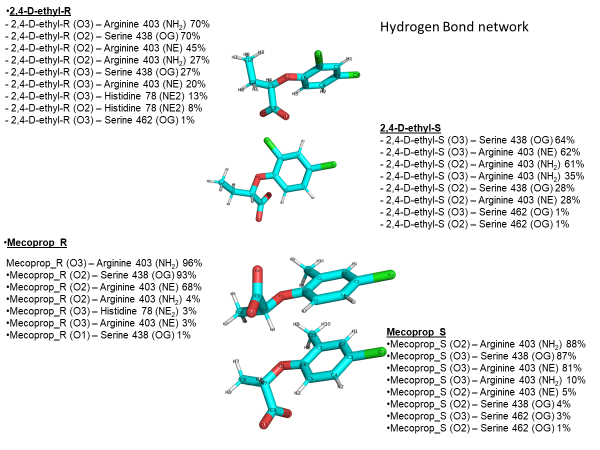

**Figure S4**. **The hydrogen bond network between the very active 2,4-D analogues and TIR1.**

### Commercially available 2,4-D analogues

**2,4-D** (CAS number 94-75-7; Sigma-Aldrich); **3,5-Me** (CAS number 5406-14-4; Sigma-Aldrich); **PHAA** (CAS number 122-59-8; Antonides CV); **PA** (CAS number 103-82-2; Fisher Scientific); **MCPA** (CAS number 94-74-6; Sigma-Aldrich); **MCPB** (CAS number 94-81-5; Sigma Aldrich); **2,4-DnB** (CAS number 94-82-6; Sigma Aldrich); **2-Cl** (CAS number 614-61-9; Fisher Scientific); **2-F** (CAS number 348-10-7; Fisher Scientific); **2-Br** (CAS number 1879-56-7; Fisher Scientific); **3-Cl** (CAS number 588-32-9; Fisher Scientific); **4-Cl** (CAS number 122-88-3; Antonides CV); **4-F** (CAS number 405-79-8; Bio-Connect); **4-Br** (CAS number 1878-91-7; Fisher Scientific); **4-I** (CAS number 1979-94-0; Fisher Scientific); **4-NO_2_** (CAS number 1798-11-4; Bio-Connect); **3-Me** (CAS number 1643-15-8; Bio-Connect); **3-Ome** (CAS number 2088-24-6; Fisher Scientific); **2,3-D** (CAS number 2976-74-1; Fisher Scientific); **3,4-D** (CAS number 588-22-7; Fisher Scientific); **2-Cl-4-Formyl** (CAS number 52268-20-9; Sigma-Aldrich); **2-Cl-4-F** (CAS number 399-41-7; Bio-Connect); **2,4-F** (CAS number 399-44-0; Bio-Connect); **2,4-Br** (CAS number 10129-78-9; Fisher Scientific); **2,4,5-T** (CAS number 93-76-5; Fisher Scientific); **2,4-DP** (CAS number 120-36-5; Sigma Aldrich); **Mecoprop** (CAS number 93-65-2; Sigma Aldrich); **2,4-DiB** (CAS number 1914-66-5; Bio-Connect); **2,4-DB** (CAS number 6956-86-1; Bio-Connect)

All commercially available compounds were tested for purity with ^1^H NMR and were >95 % pure, except for compound **4-I** which was circa 90% pure.

### Chemical synthesis

General procedure for the synthesis of 2,4-D analogues

The corresponding phenol of the analogue was charged to a roundbottom flask together with potassium hydroxide, bromo-or iodoacetic acid, and water. The reaction mixture was heated at 100 °C for several hours, typically five to six hours. Water was added during the reaction if the reaction mixture dried up or the reaction mixture was diluted before work-up. The reaction mixture was acidified to pH ≤ 1 and extracted with Et_2_O. The organic layer was washed with brine, dried over Na_2_SO_4_, filtered and concentrated *in vacuo*. The crude was separated using flash column chromatography. A detailed experimental procedure per analogue can be found in the SI.

**2-I**

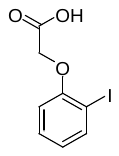
2-iodophenol (580 mg; 2.6 mmol), potassium hydroxide (1.008 g; 18.0 mmol), iodoacetic acid (1.469 g; 7.9 mmol) and water (50 mL) were charged to a roundbottom flask. The reaction mixture was heated at 100 °C for several hours. Water (20 mL) was added during the reaction, because some evaporated. The reaction mixture was diluted with water (20 mL), acidified with 6 M HCl to pH ≤ 1 and extracted with Et_2_O (3 x 20 mL). The organic layer was washed with brine (3 x 20 mL). The organic layer was dried over Na_2_SO_4_, filtered and concentrated *in vacuo* to yield a yellow oil. The crude was dry loaded and column chromatography (70 % PE ; 30 % EtOAc; 1 % acetic acid) yielded **2-I** (432 mg; 1.6 mmol; 59 %).

^1^H NMR (600 MHz, CDCl3) δ 7.81 (dd, J = 7.8, 1.6 Hz, 1H), 7.33 (ddd, J = 8.2, 7.4, 1.6 Hz, 1H), 6.85 – 6.75 (m, 2H), 4.72 (s, 2H).

^13^C NMR (600 MHz, CDCl3) δ 169.91, 155.75, 139.83, 129.73, 124.27, 112.63, 86.48, 65.90.

ESI HRMS for C_8_H_7_IO_3_: m/z [M – H]^-^ ; calcd: 276,9362; found: 276,9349.

**2,5-D**

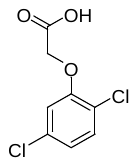
2,5-dichlorophenol (0.296 g; 1.8 mmol), potassium hydroxide (3.57 g; 63.6 mmol), bromoacetic acid (3.68 g; 26.5 mmol) and water (50 mL) were charged to a roundbottom flask. The reaction mixture was heated at 100 °C for several hours (3) until it was concentrated. The reaction mixture was diluted with water (50 mL), acidified with 6 M HCl to pH ≤ 1 and extracted with Et_2_O (3 x 30 mL). The organic layer was washed with water (6 x 15 mL). The organic layer was dried over Na_2_SO_4_, filtered and concentrated in vacuo. The Et_2_O was washed with brine and dried over Na_2_SO_4_, filtered and concentrated *in vacuo*. Column chromatography (70 % PE ; 30 % EtOAc; 1 % acetic acid) yielded **2,5-D** (0.179 g; 0.84 mmol; 47 %).

^1^H NMR (600 MHz, DMSO) δ 7.53 (d, J = 8.5 Hz, 1H), 7.23 (d, J = 2.3 Hz, 1H), 7.11 (dd, J = 8.5, 2.3 Hz, 1H), 4.94 (s, 2H).

^13^C NMR (600 MHz, DMSO) δ 169.91, 154.33, 132.67, 131.50, 122.16, 120.66, 114.55, 65.68.

ESI HRMS for C_8_H_6_Cl_2_O_3_: m/z [M – H]^-^ ; calcd: 218,9616; found: 218,9609.

**2,6-D**

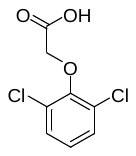
2,6-dichlorophenol (0.315 g; 1.9 mmol), potassium hydroxide (3.7 g; 65.9 mmol), bromoacetic acid (3.86 g; 27.8 mmol) and water (50 mL) were charged to a roundbottom flask. The reaction mixture was heated at 100 °C for several hours (6). The reaction mixture was diluted with water (25 mL), acidified with 6 M HCl to pH ≤ 1 and extracted with Et_2_O (3 x 25 mL). The organic layer was washed with brine (3 x 25 mL). The organic layer was dried over Na_2_SO_4_, filtered and concentrated *in vacuo*. The organic layer was dried over Na_2_SO_4_, filtered and concentrated in vacuo. Column chromatography (70 % PE ; 30 % EtOAc; 1 % acetic acid) yielded **2,6-D** (0.069 g; 0.31 mmol; 16% ).

^1^H NMR (600 MHz, CDCl3) δ 7.34 (d, J = 8.1 Hz, 2H), 7.09 (t, J = 8.1 Hz, 1H), 4.69 (s, 2H).

^13^C NMR (600 MHz, DMSO) δ 169.30, 150.32, 129.86, 128.71, 126.87, 69.45.

ESI HRMS for C_8_H_6_Cl_2_O_3_: m/z [M – H]^-^ ; calcd: 218,9616; found: 218,9603.

**3,5-D**

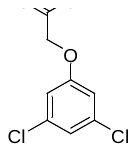
3,5-dichlorophenol (0.303 g; 1.9 mmol), potassium hydroxide (3.65 g; 65.1 mmol), bromoacetic acid (3.91 g; 28.1 mmol) and water (50 mL) were charged to a roundbottom flask. The reaction mixture was heated at 100 °C for several hours (5.5). The reaction mixture was diluted with water (25 mL), acidified with 6 M HCl to pH ≤ 1 and extracted with Et_2_O (3 x 25 mL). The organic layer was washed with brine (3 x 25 mL). The organic layer was dried over Na_2_SO_4_, filtered and concentrated *in vacuo*. The organic layer was dried over Na_2_SO_4_, filtered and concentrated *in vacuo*. Column chromatography (70 % PE ; 30 % EtOAc; 1 % acetic acid) yielded **3,5-D** (0.199 g; 0.9 mmol; 48% ).

^1^H NMR (600 MHz, CDCl3) δ 7.04 (s, 1H), 6.83 (s, 2H), 4.68 (s, 2H).

^13^C NMR (600 MHz, DMSO) δ 170.05, 159.70, 134.98, 121.23, 114.50, 65.37.

ESI HRMS for C_8_H_6_Cl_2_O_3_: m/z [M – H]^-^ ; calcd: 218,9616; found: 218,9608.

**2-F-4-Br**

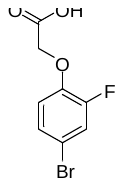
2-fluoro-4-bromophenol (0.508 g; 2.7 mmol), potassium hydroxide (1.051 g; 18.7 mmol), iodoacetic acid (1.441 g; 7.7 mmol) and water (50 mL) were charged to a roundbottom flask. The reaction mixture was heated at 100 °C for several hours. Water (20 mL) was added during the reaction and before work-up (10mL). The reaction mixture was acidified with 6 M HCl to pH ≤ 1 and extracted with Et_2_O (3 x 25 mL). The organic layer was washed with brine (3 x 25 mL). The organic layer was dried over Na_2_SO_4_, filtered and concentrated *in vacuo*. Column chromatography (70 % PE ; 30 % EtOAc; 1 % acetic acid) yielded **2-F-4-Br** (0.441 g; 1.8 mmol; 67 %).

^1^H NMR (600 MHz, DMSO) δ 13.14 (s, 1H), 7.55 (dd, J = 10.9, 2.4 Hz, 1H), 7.32 (ddd, J = 8.8, 2.4, 1.5 Hz, 1H), 7.07 (t, J = 9.0 Hz, 1H), 4.80 (s, 2H).

^13^C NMR (600 MHz, DMSO) δ 170.05, 152.80, 151.15, 145.83, 145.76, 127.92, 127.89, 119.90, 119.75, 117.02, 117.01, 112.00, 111.95, 65.55.

^19^F NMR (565 MHz, DMSO) δ -131.29 (t, J = 10.1 Hz).

ESI HRMS for C_8_H_6_BrFO_3_: m/z [M – H]^-^ ; calcd: 246,9406; found: 246,9394.

**2-F-4-Cl**

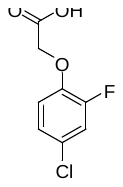
2-fluoro-4-chlorophenol (491 mg; 3.4 mmol), potassium hydroxide (1.34 g; 24.0 mmol), iodoacetic acid (1.90 g; 10.2 mmol) and water (50 mL) were charged to a roundbottom flask. The reaction mixture was heated at 100 °C for several hours. Water (20 mL) was added during the reaction, because some evaporated. The reaction mixture was acidified with 6 M HCl to pH 3 and a white solid was observed. The aqueous layer was extracted with Et_2_O (3 x 30 mL). The organic layer was washed with brine (3 x 30 mL) while the white solid remained in the organic layer. The organic layer was dried over Na_2_SO_4_, filtered and concentrated *in vacuo*. The crude was again dissolved in Et_2_O and washed with water (3 x 30 mL). Column chromatography (70 % PE ; 30 % EtOAc; 1 % acetic acid) yielded **2-F-4-Cl** (78 mg; 0.4 mmol; 20 %).

^1^H NMR (600 MHz, CDCl3) δ 7.15 (dd, J = 10.8, 2.5 Hz, 1H), 7.06 (ddd, J = 8.8, 2.5, 1.6 Hz, 1H), 6.91 (t, J = 8.8 Hz, 1H), 4.74 (s, 2H).

^13^C NMR (600 MHz, CDCl3) δ 172.65, 153.43, 151.77, 144.37, 127.57, 124.51, 124.48, 117.66, 117.51, 117.18, 66.38.

^19^F NMR (400 MHz, CDCl3) δ -129.94 (t, J = 9.7 Hz).

ESI HRMS for C_8_H_6_ClFO_3_: m/z [M – H]^-^ ; calcd: 202,9911; found: 202,9906.

**2-NO_2_-4-Cl**

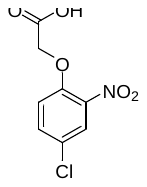
2-nitro-4-chlorophenol (0.311 g; 1.8 mmol), potassium hydroxide (0.714 g; 12.7 mmol), iodoacetic acid (0.967 g; 5.2 mmol) and water (50 mL) were charged to a roundbottom flask. The reaction mixture was heated at 100 °C for several hours. Water was added during the reaction (3 x 10 mL) and before work-up (10mL). The reaction mixture was acidified with 6 M HCl to pH ≤ 1 and extracted with Et_2_O (3 x 25 mL). The organic layer was washed with brine (4 x 20 mL). The organic layer was dried over Na_2_SO_4_, filtered and concentrated *in vacuo*. Column chromatography (70 % PE ; 30 % EtOAc; 1 % acetic acid; switched to 100 % EtOAc; 1 % acetic acid) yielded **2-NO_2_-4-Cl** (0.093 g; 0.4 mmol; 22 %).

^1^H NMR (600 MHz, DMSO) δ 8.04 (d, J = 2.7 Hz, 1H), 7.70 (dd, J = 9.1, 2.7 Hz, 1H), 7.32 (d, J = 9.1 Hz, 1H), 4.93 (s, 2H).

^13^C NMR (600 MHz, DMSO) δ 169.61, 149.77, 140.49, 134.03, 124.90, 124.87, 117.47, 66.08.

ESI HRMS for C_8_H_6_ClNO_5_: m/z [M – H]^-^ ; calcd: 229,9856; found: 229,9851.

**2,3,4-T**

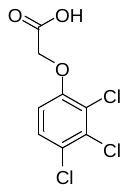
2,3,4-trichlorophenol (0.051 g; 0.3 mmol), potassium hydroxide (0.099 g; 1.8 mmol), iodoacetic acid (0.174 g; 0.9 mmol) and water (10 mL) were charged to a roundbottom flask. The reaction mixture was heated at 100 °C for several hours. Water was added during the reaction (4 x 10 mL). The reaction mixture was acidified with 6 M HCl to pH ≤ 1 and extracted with Et_2_O (3 x 20 mL). The organic layer was washed with brine (2 x 20 mL). The organic layer was dried over Na_2_SO_4_, filtered and concentrated *in vacuo*. Column chromatography (70 % PE ; 30 % EtOAc; 1 % acetic acid) yielded **2,3,4-T** (0.031 g; 0.1 mmol; 47 %). NMR analysis demonstrated that the compound, most likely, contained grease impurities.

^1^H NMR (600 MHz, MeOD) δ 7.42 (d, J = 9.1 Hz, 1H), 6.98 (d, J = 9.1 Hz, 1H), 4.81 (s, 2H).

^13^C NMR (600 MHz, MeOD) δ 176.58/170.09, 153.88, 131.69, 127.86, 125.24, 123.01, 112.15, 70.18/68.88/65.47/63.01.

ESI HRMS for C_8_H_5_Cl_3_O_3_: m/z [M – H]^-^ ; calcd: 252,9226; found: 252,9211.

**2,4,6-T**

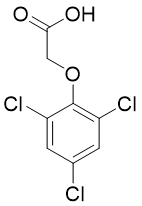
2,4,6-trichlorophenol (0.346 g; 1.8 mmol), potassium hydroxide (3.5 g; 62.4 mmol), bromoacetic acid (3.6 g; 25.9 mmol) and water (50 mL) were charged to a roundbottom flask. The reaction mixture was heated at 100 °C overnight. Water was added before the work-up (50mL). The reaction mixture was acidified with 6 M HCl to pH ≤ 1 and extracted with Et_2_O (3 x 25 mL). The organic layer was washed with brine (3 x 25 mL). The organic layer was dried over Na_2_SO_4_, filtered and concentrated *in vacuo*. Column chromatography (70 % PE ; 30 % EtOAc; 1 % acetic acid) yielded **2,4,6-T** (0.055 g; 0.2 mmol; 12 %).

^1^H NMR (600 MHz, DMSO) δ 7.72 (s, 2H), 4.59 (s, 2H).

^13^C NMR (600 MHz, DMSO) δ 169.19, 149.71, 129.67, 129.57, 129.43, 69.50.

ESI HRMS for C_8_H_5_Cl_3_O_3_: m/z [M – H]^-^ ; calcd: 252,9226; found: 252,9211.

**2,3,4-F**

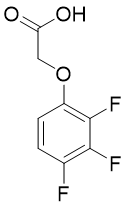
2,3,4-trifluorophenol (300 mg; 2.0 mmol), potassium hydroxide (0.928 g; 16.5 mmol), iodoacetic acid (1.139 g; 6.1 mmol) and water (40 mL) were charged to a roundbottom flask. The reaction mixture was heated at 100 °C for several hours. Water (10 mL) was added during the reaction, because some evaporated. The reaction mixture was acidified with 6 M HCl to pH ≤ 1 and extracted with Et_2_O (3 x 20 mL). The organic layer was washed with brine (3 x 20 mL). The organic layer was dried over Na_2_SO_4_, filtered and concentrated *in vacuo*. The crude was dry loaded and column chromatography (70 % PE ; 30 % EtOAc; 1 % acetic acid) yielded **2,3,4-F** (106 mg; 0.51 mmol; 25 %).

^1^H NMR (600 MHz, MeOD) δ 6.91 (tdd, J = 10.2, 8.3, 2.5 Hz, 1H), 6.76 (tdd, J = 9.1, 4.5, 2.5 Hz, 1H), 4.80 (s, 3H).

^13^C NMR (600 MHz, MeOD) δ 170.32, 110.24, 110.21, 110.11, 110.08, 108.97, 108.92, 65.82.

^19^F NMR (400 MHz, CDCl3) δ -141.53 (m), -152.26 (m), -157.15 – -157.33 (m).

ESI HRMS for C_8_H_5_F_3_O_3_: m/z [M – H]^-^ ; calcd: 205,0113; found: 205,0105.

**2,4,6-F**

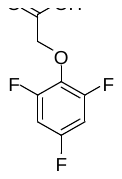
2,4,6-trifluorophenol (280 mg; 1.9 mmol), potassium hydroxide (1.1 g; 19.6 mmol), iodoacetic acid (1.306 g; 7.0 mmol) and water (30 mL) were charged to a roundbottom flask. The reaction mixture was heated at 100 °C for several hours. Water (10 mL) was added during the reaction, because some evaporated. The reaction mixture was acidified with 6 M HCl to pH ≤ 1 and extracted with Et_2_O (3 x 20 mL). The organic layer was washed with brine (3 x 20 mL). The organic layer was dried over Na_2_SO_4_, filtered and concentrated *in vacuo* to yield a yellow oil. Column chromatography (70 % PE ; 30 % EtOAc; 1 % acetic acid) yielded **2,4,6-F** (78 mg; 0.4 mmol; 20 %).

^1^H NMR (400 MHz, CDCl3) δ 6.78 – 6.65 (m, 1H), 4.75 (d, J = 0.9 Hz, 2H).

^13^C NMR (400 MHz, CDCl3) δ 172.43, 101.28, 101.01, 100.74, 69.48.

^19^F NMR (400 MHz, CDCl3) δ -113.28 – -113.48 (m), -124.61 (dd, J = 8.6, 2.9 Hz).

ESI HRMS for C_8_H_5_F_3_O_3_: m/z [M – H]^-^ ; calcd: 205,0113; found: 205,0101.

**2,4-DnP**

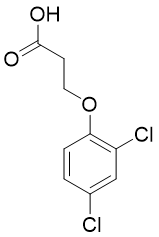
2,4-dichlorophenol (495 mg; 3.0 mmol), potassium hydroxide (1.187 g; 21.2 mmol), bromopropanoic acid (1.354 g; 8.9 mmol) and water (50 mL) were charged to a roundbottom flask. The reaction mixture was heated at 100 °C for several hours. The reaction mixture was diluted with water (20 mL), acidified with 6 M HCl to pH ≤ 1 and extracted with Et_2_O (3 x 20 mL). The organic layer was washed with brine (3 x 20 mL). The organic layer was dried over Na_2_SO_4_, filtered and concentrated *in vacuo* to yield a yellow oil. Column chromatography (70 % PE ; 30 % EtOAc; 1 % acetic acid) yielded **2,4-DnP** (136 mg; 0.6 mmol; 20 %).

^1^H NMR (600 MHz, CDCl3) δ 7.36 (d, J = 2.5 Hz, 1H), 7.18 (dd, J = 8.7, 2.5 Hz, 1H), 6.88 (d, J = 8.8 Hz, 1H), 4.29 (t, J = 6.3 Hz, 2H), 2.92 (t, J = 6.3 Hz, 2H).

^13^C NMR (600 MHz, CDCl3) δ 175.97, 152.92, 130.13, 127.63, 126.47, 124.16, 114.73, 64.79, 34.11.

ESI HRMS for C_9_H_8_Cl_2_O_3_: m/z [M – H]^-^ ; calcd: 232,9772; found: 292,9764.

### Copies of NMR spectra

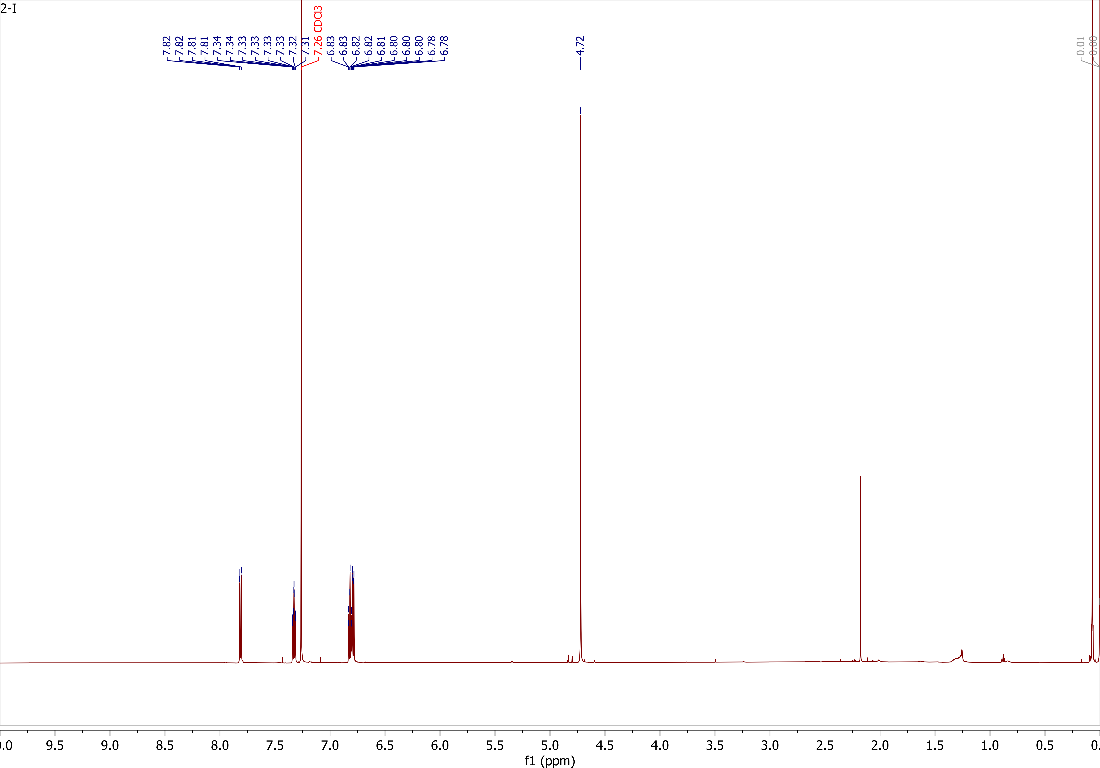

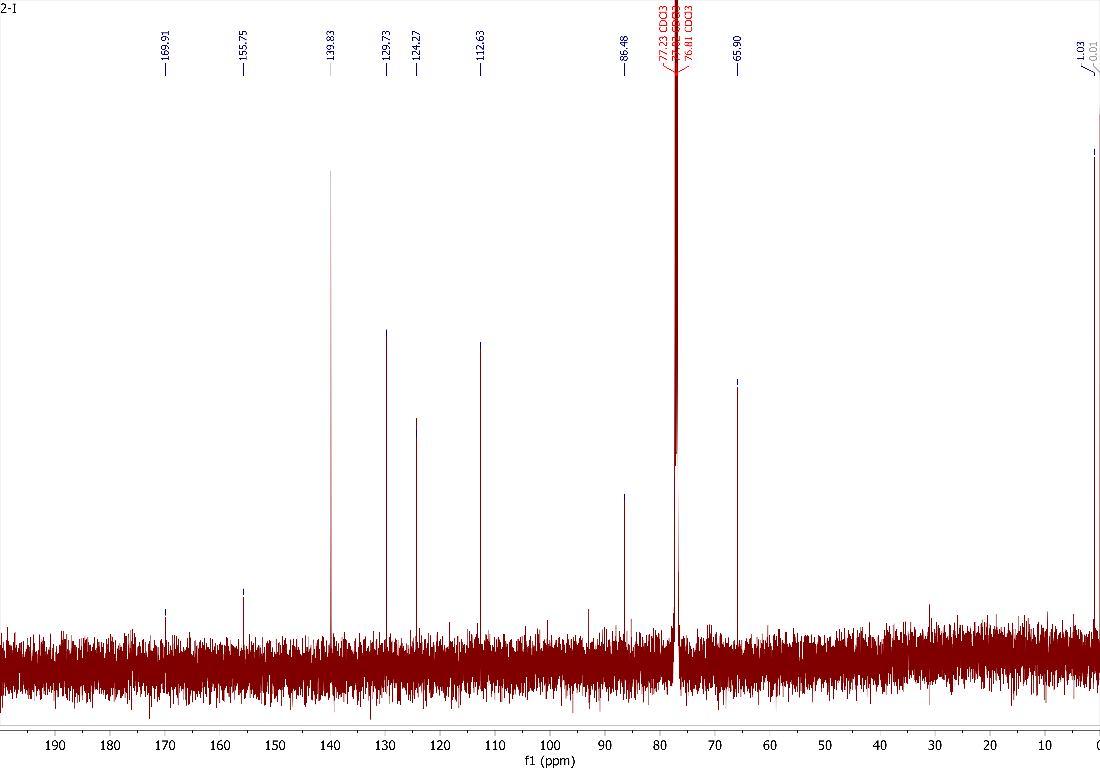

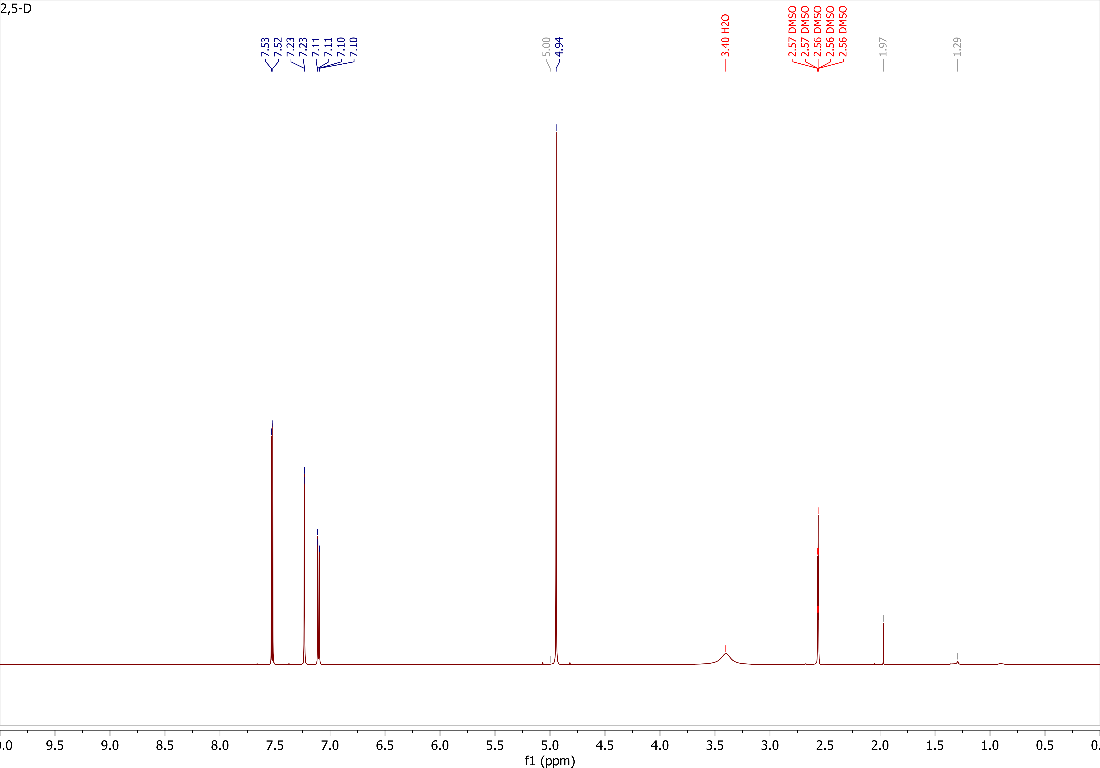

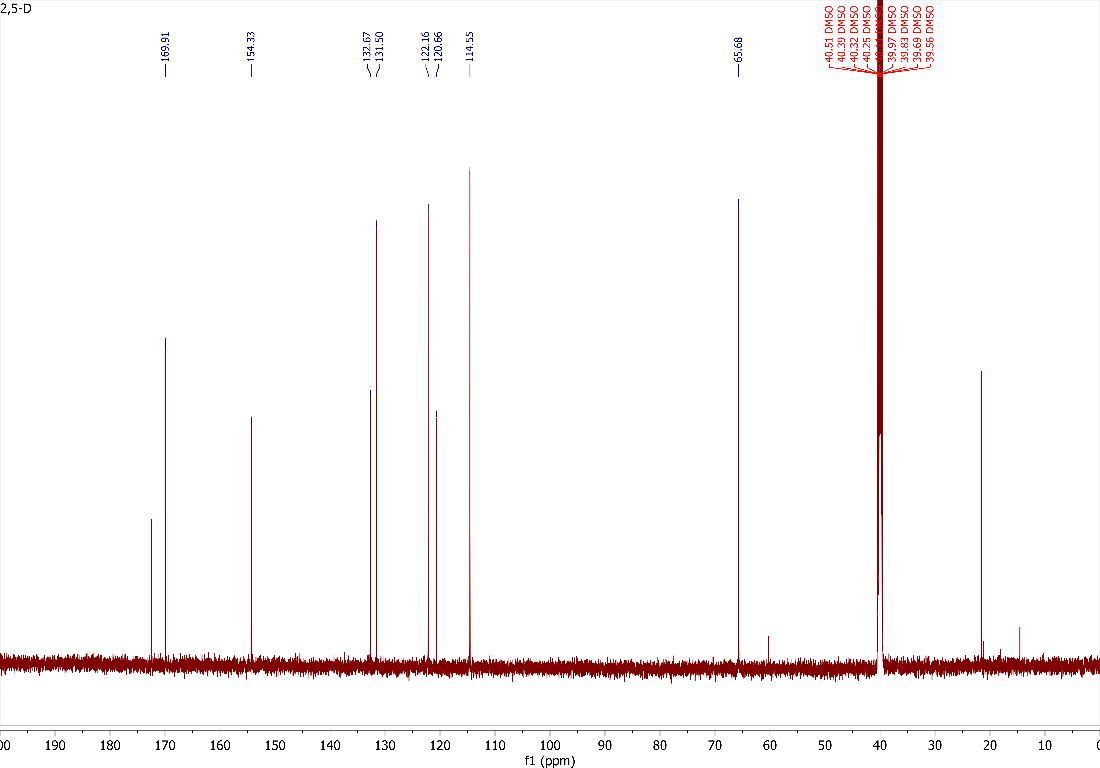

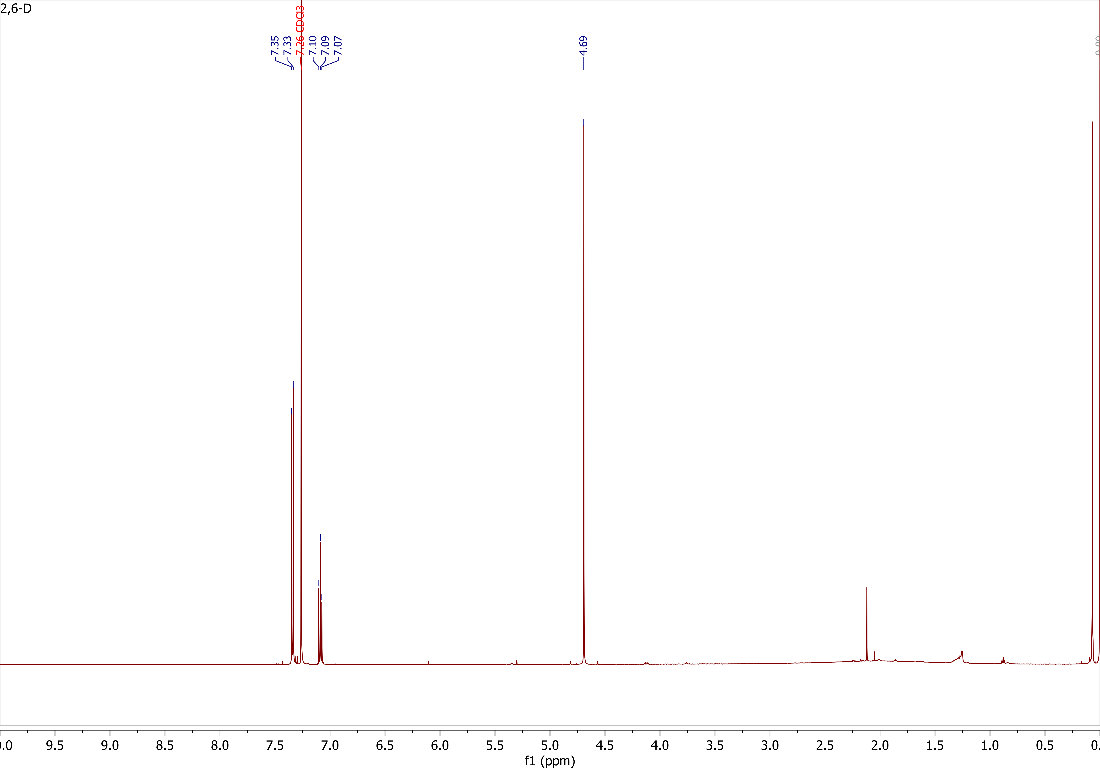

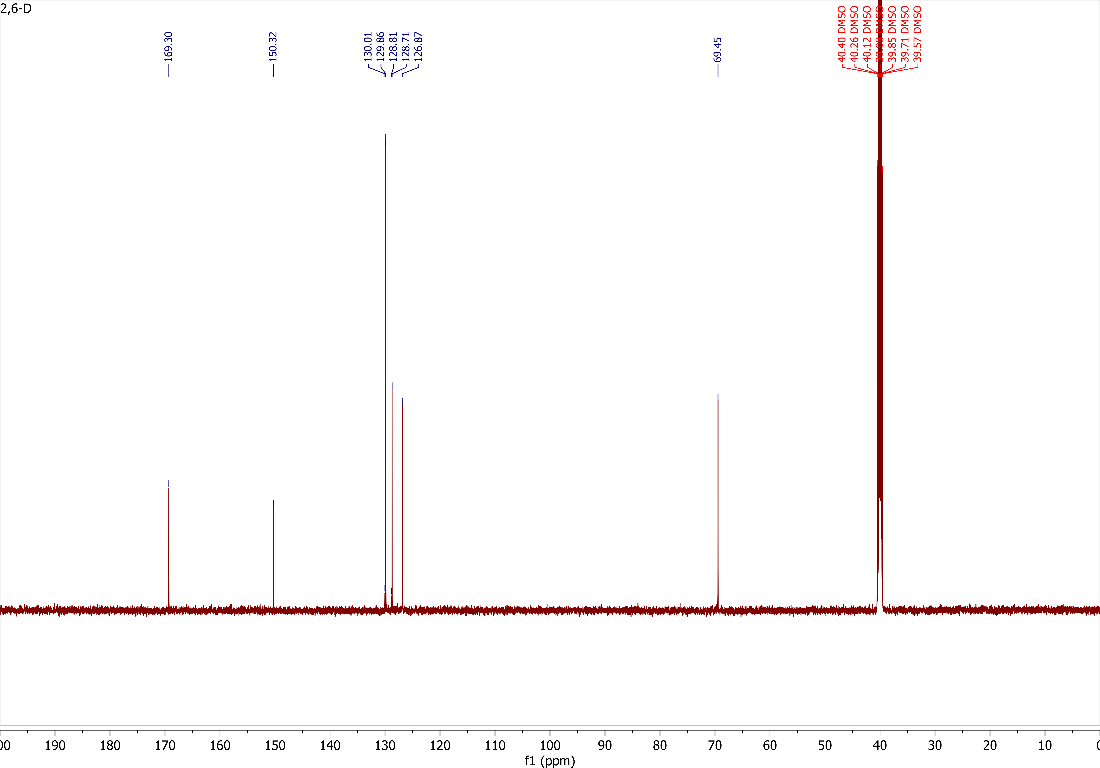

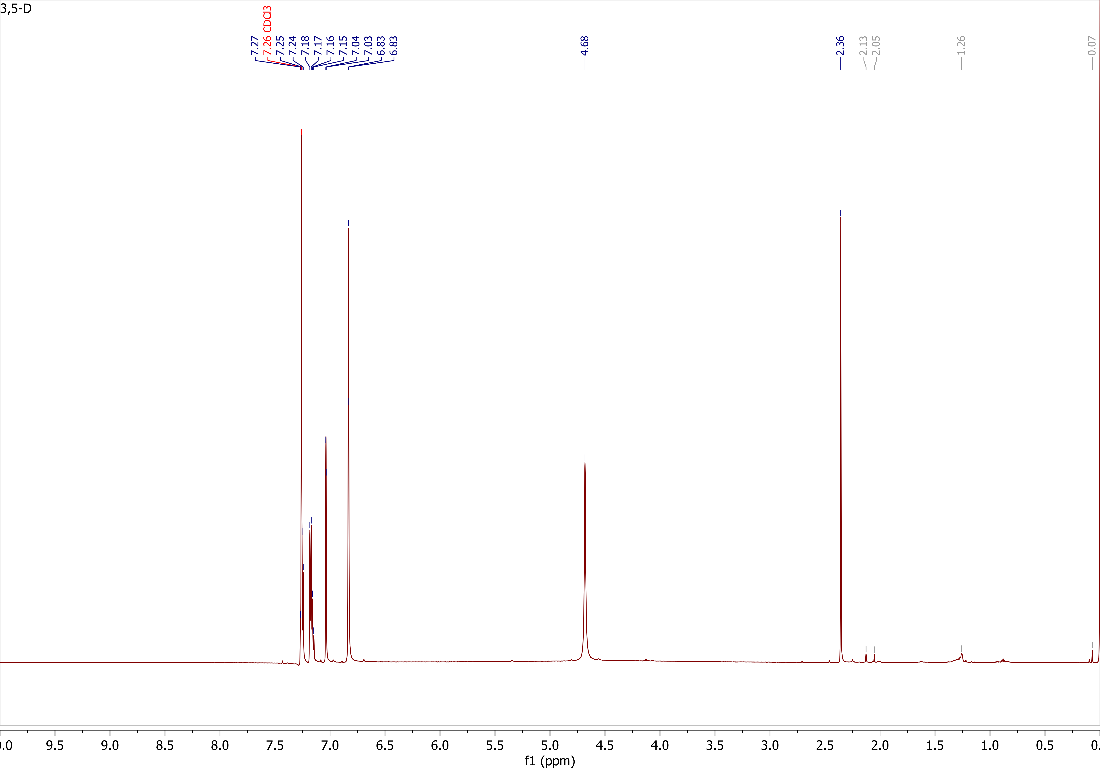

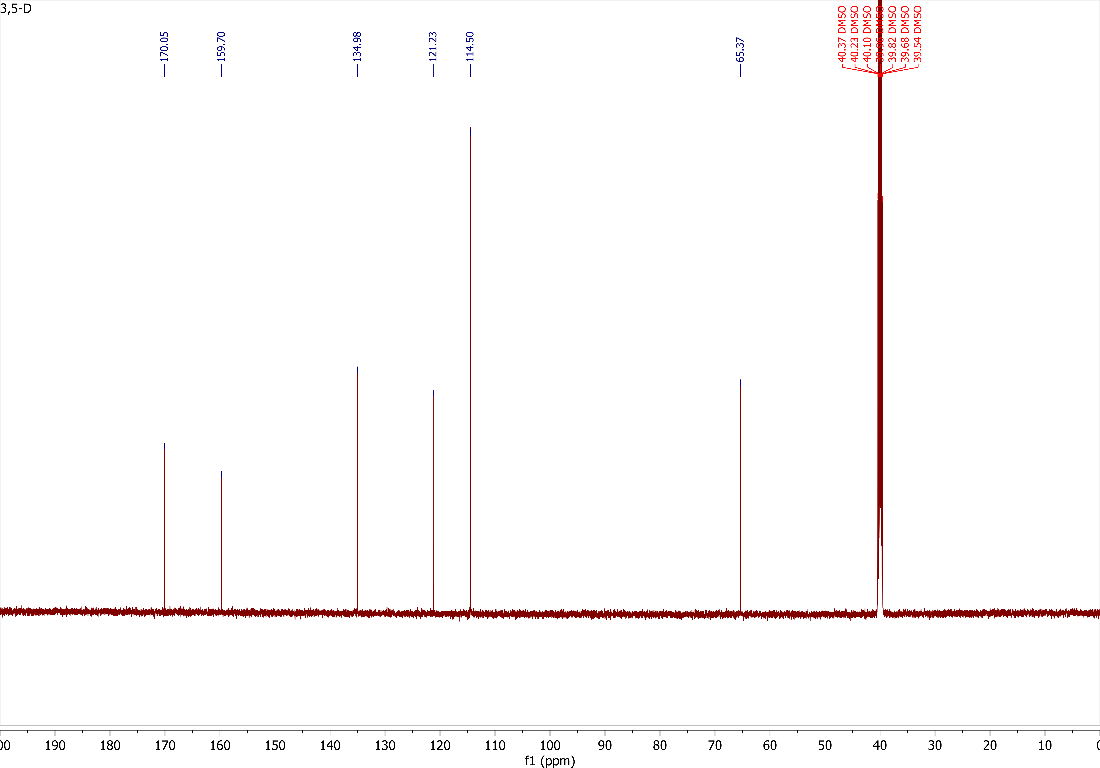

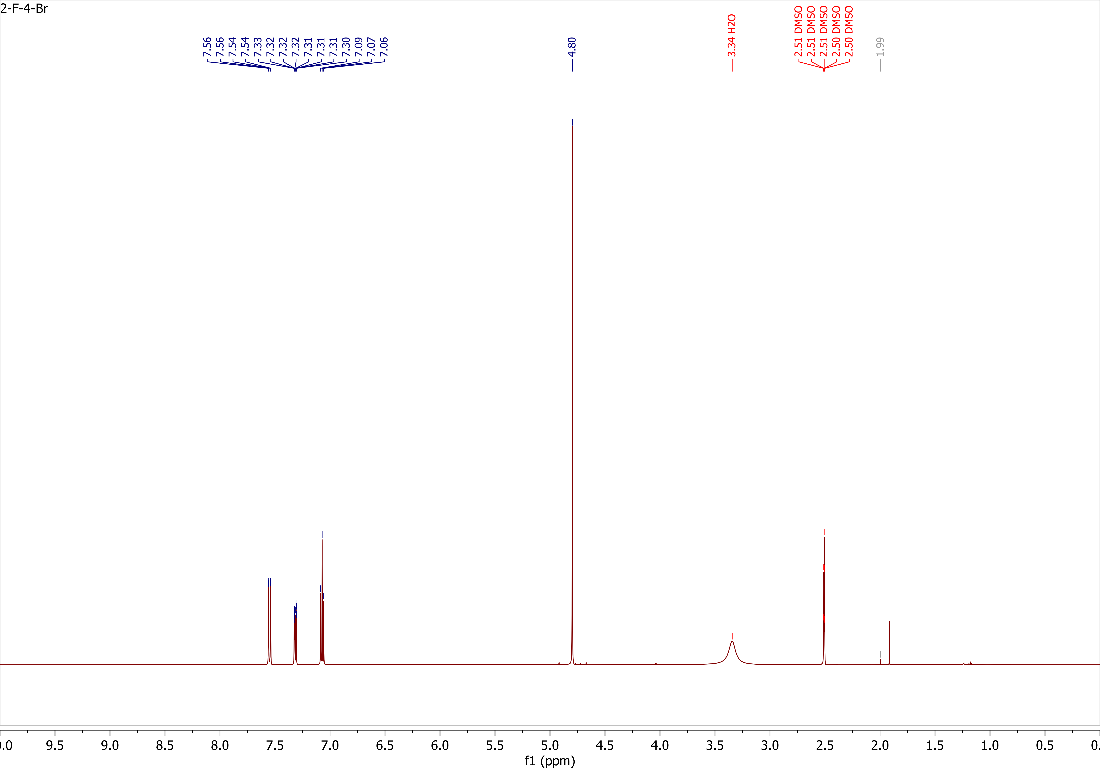

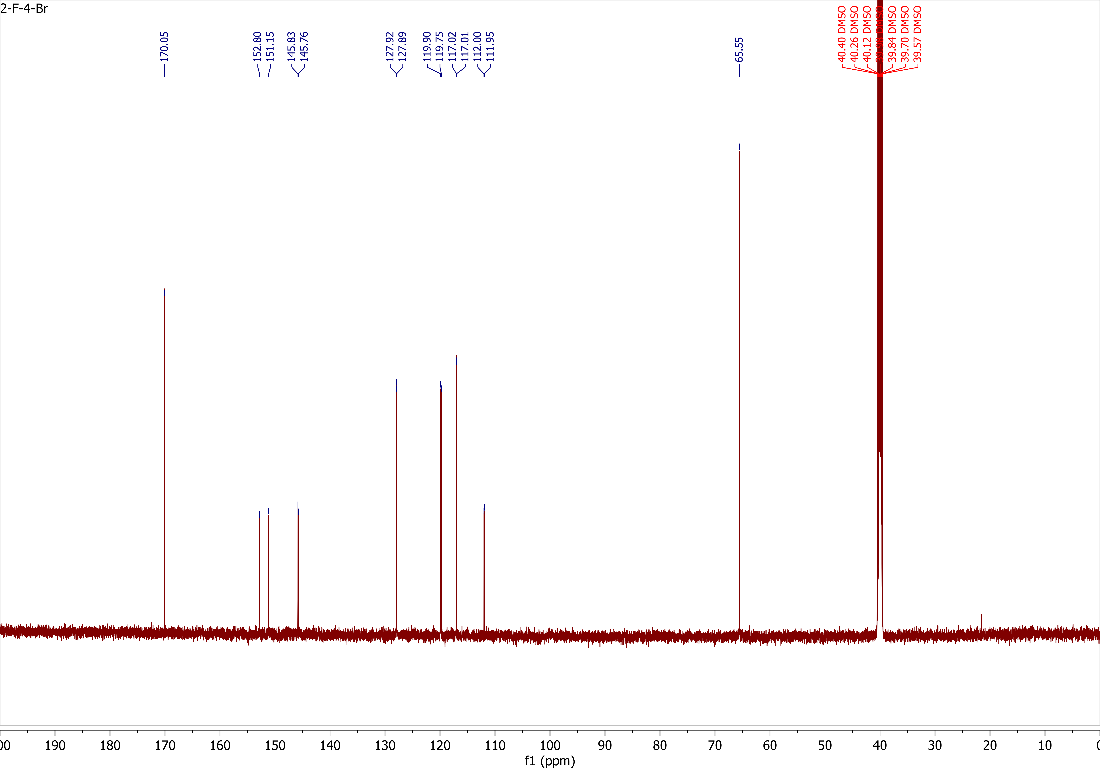

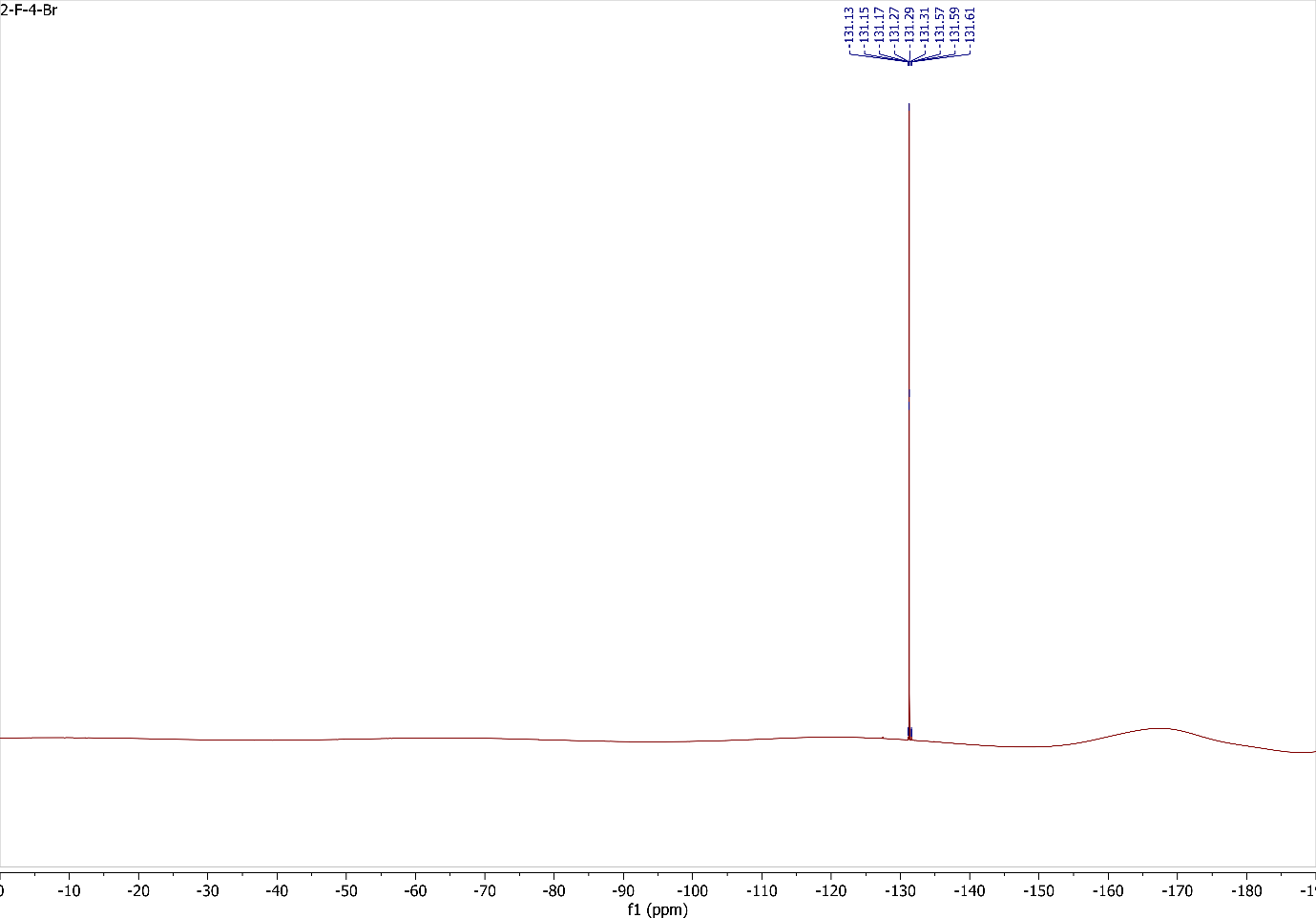

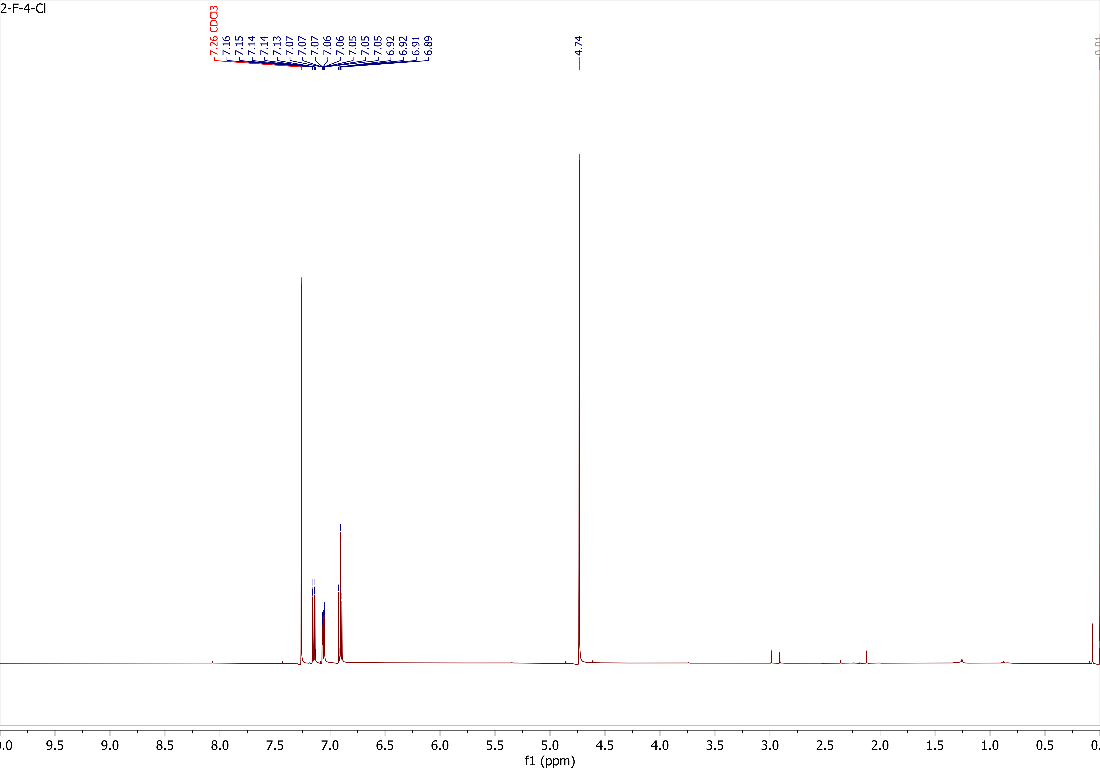

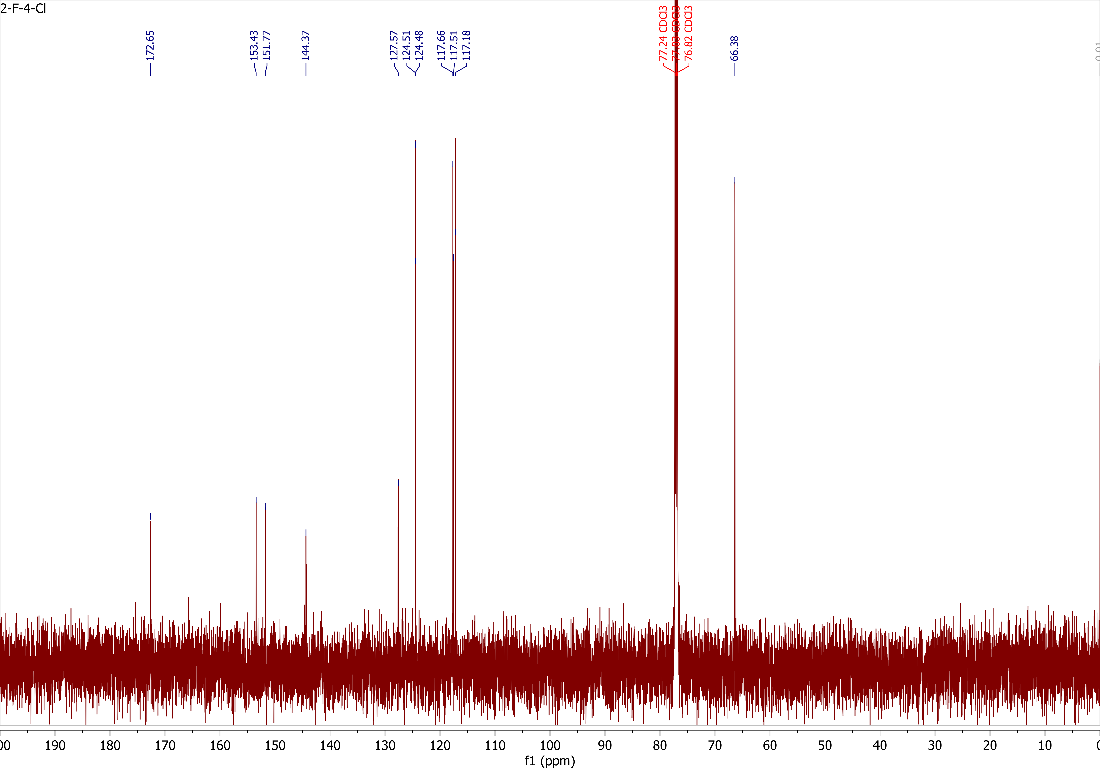

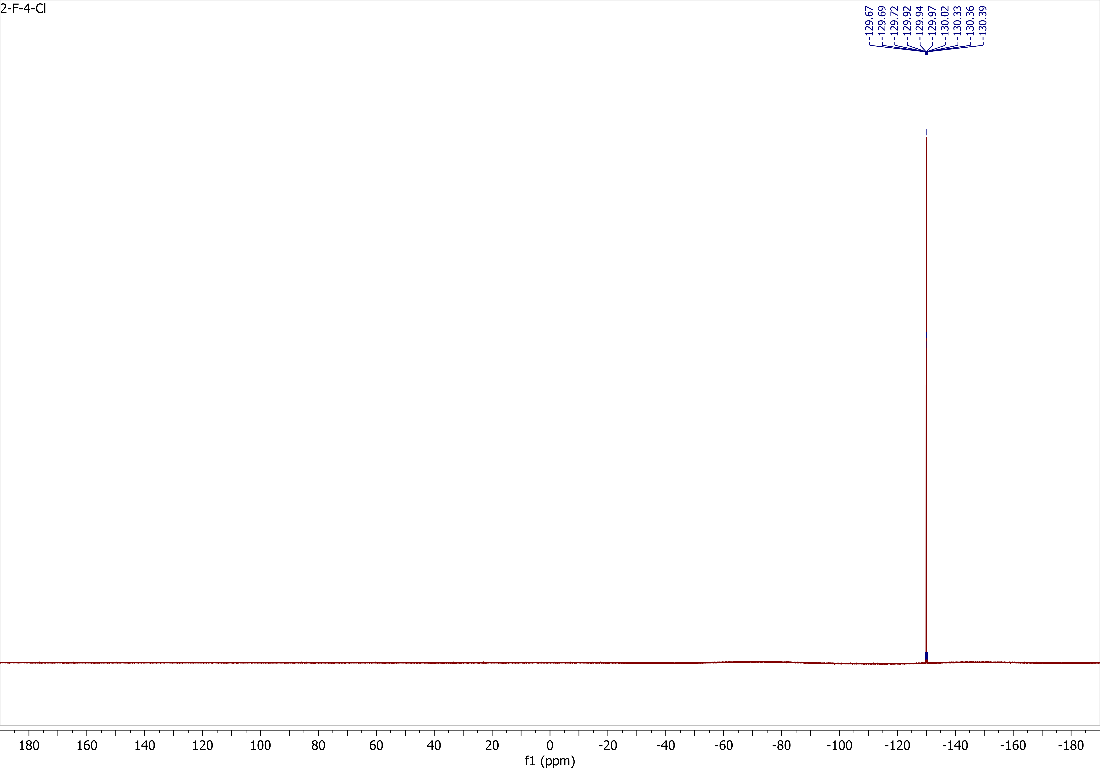

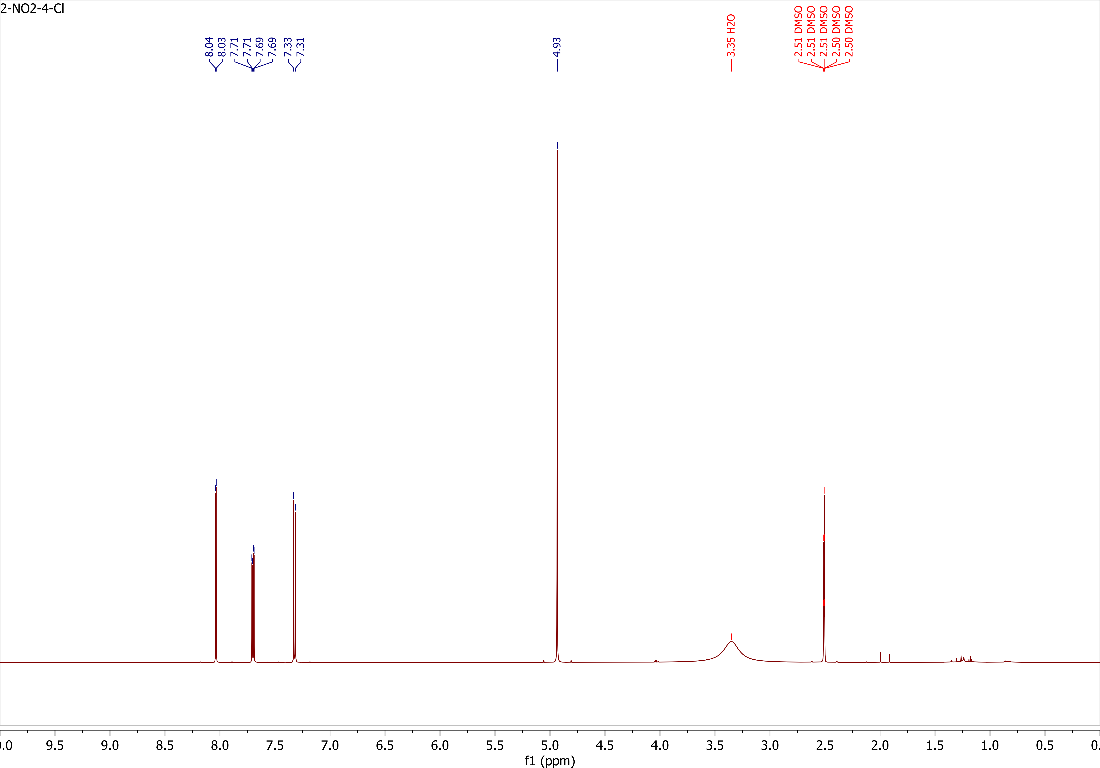

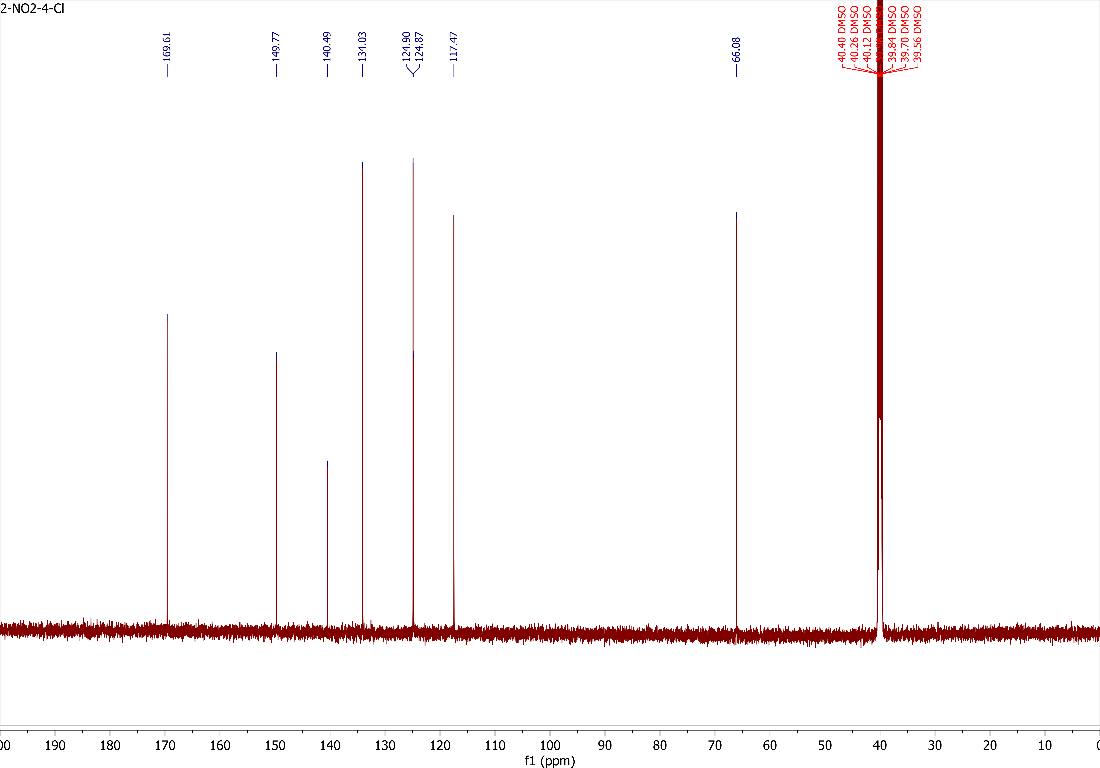

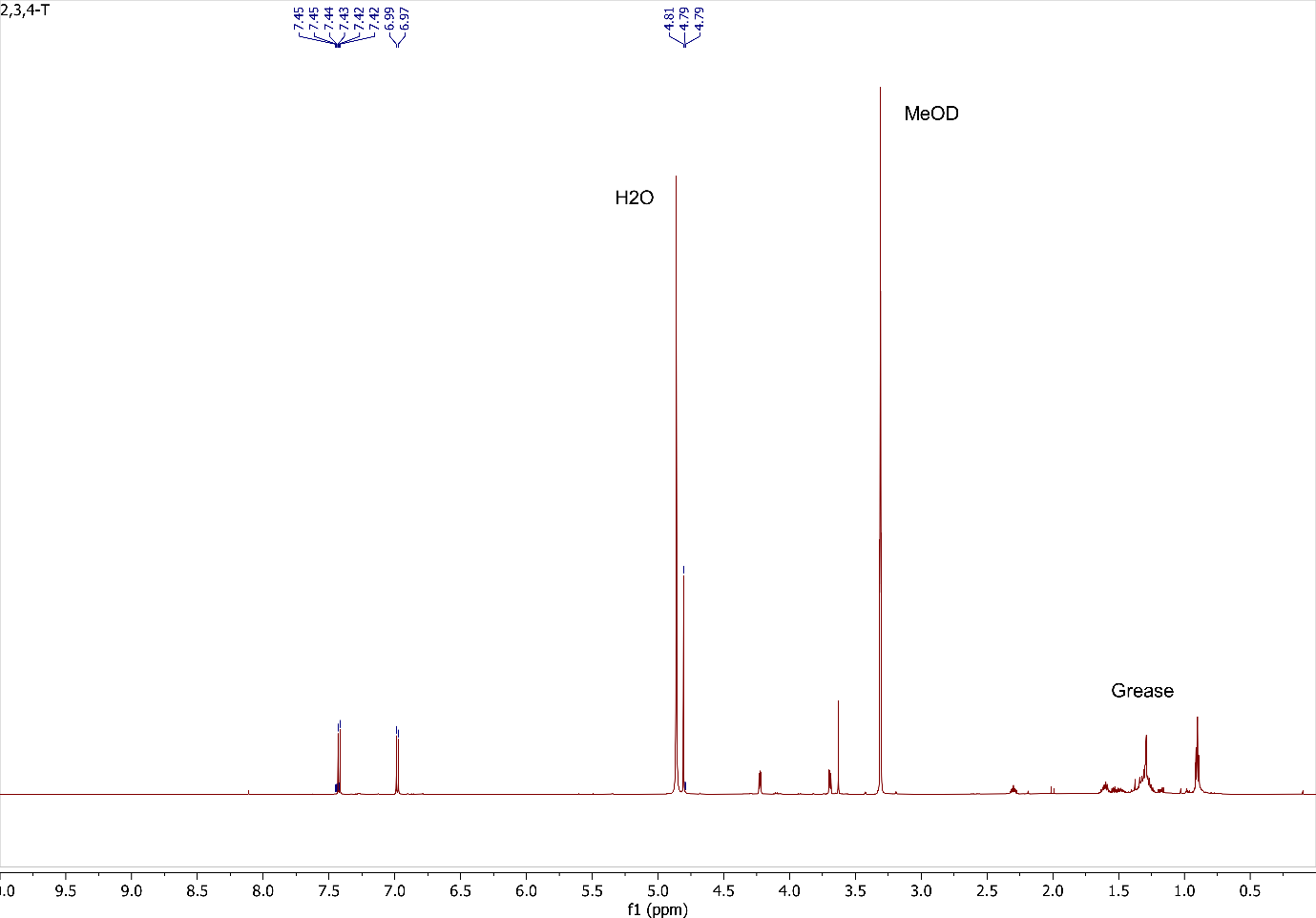

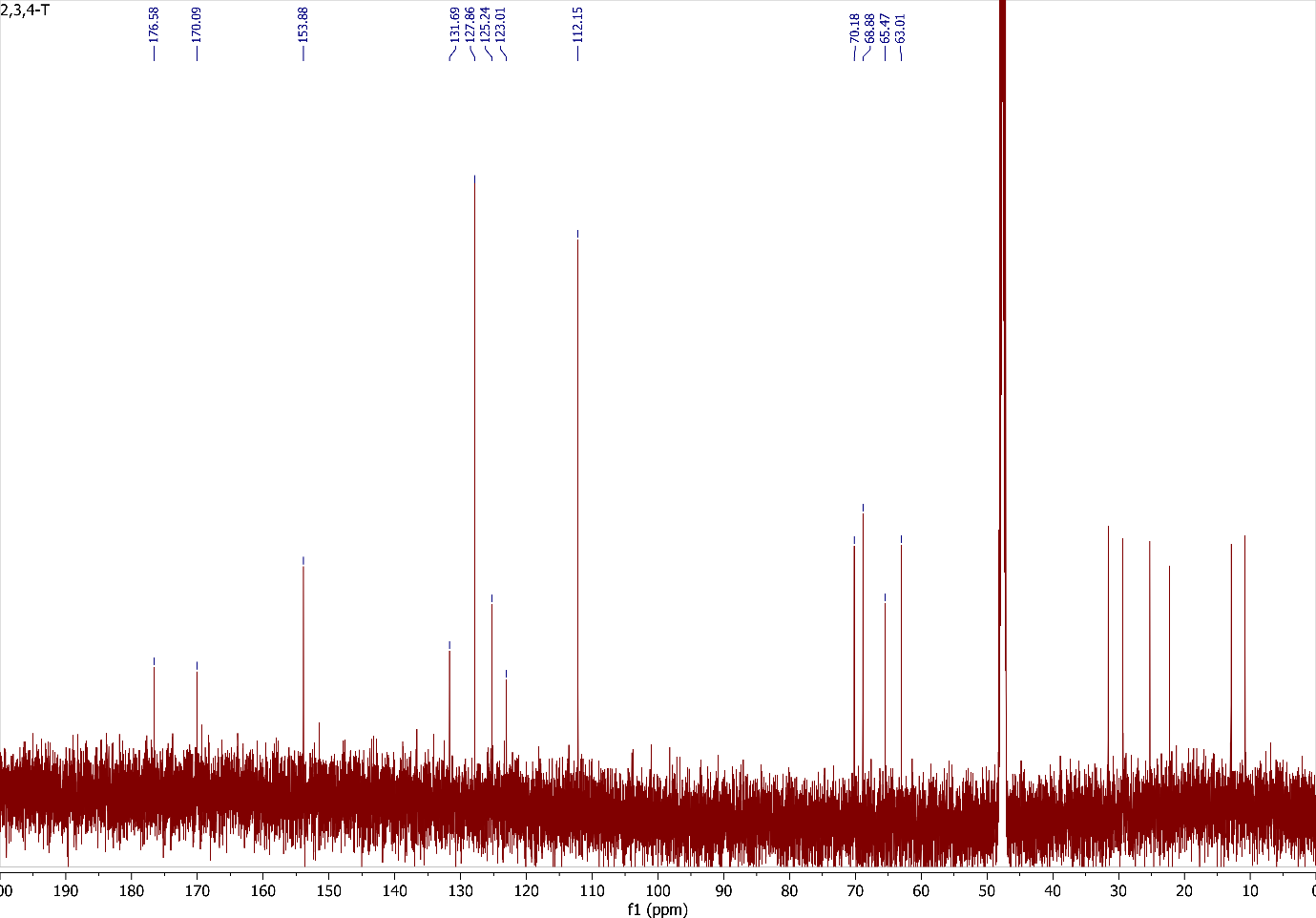

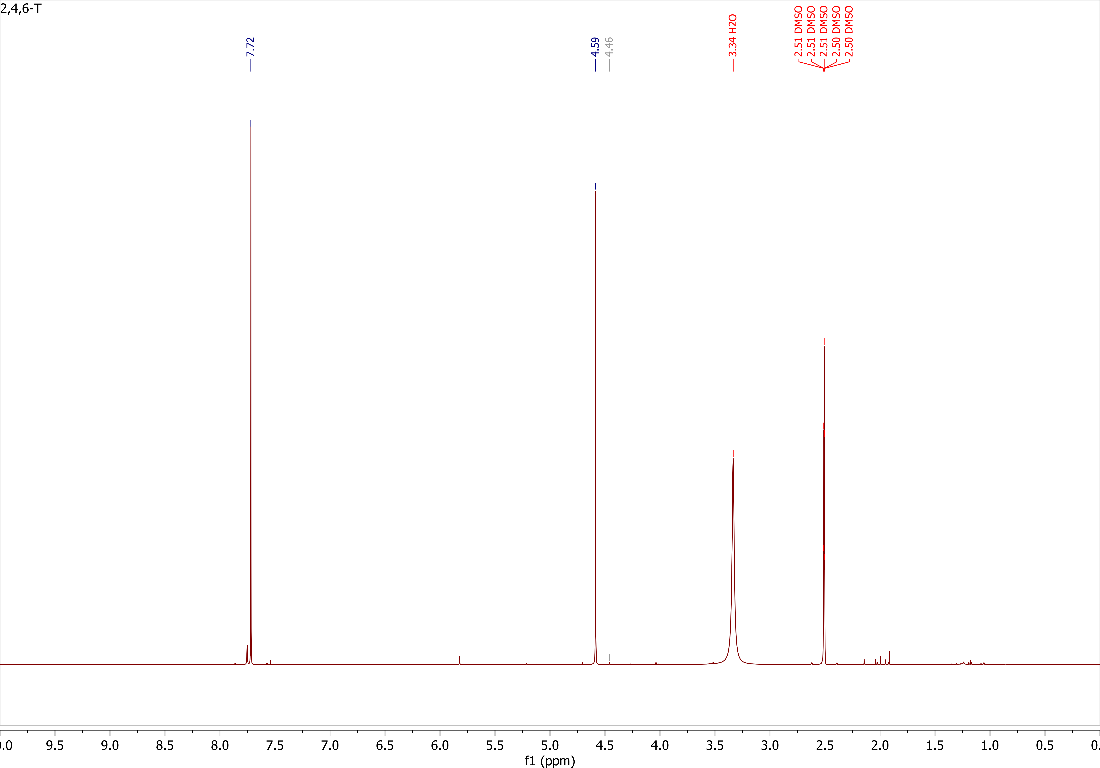

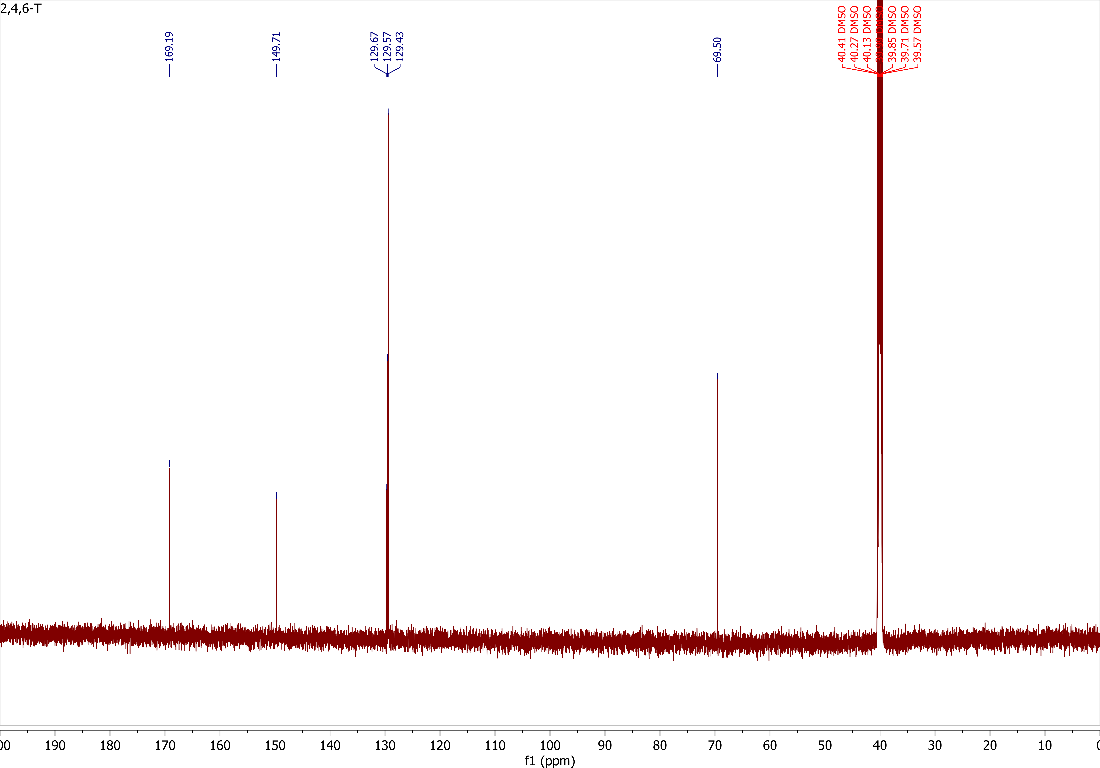

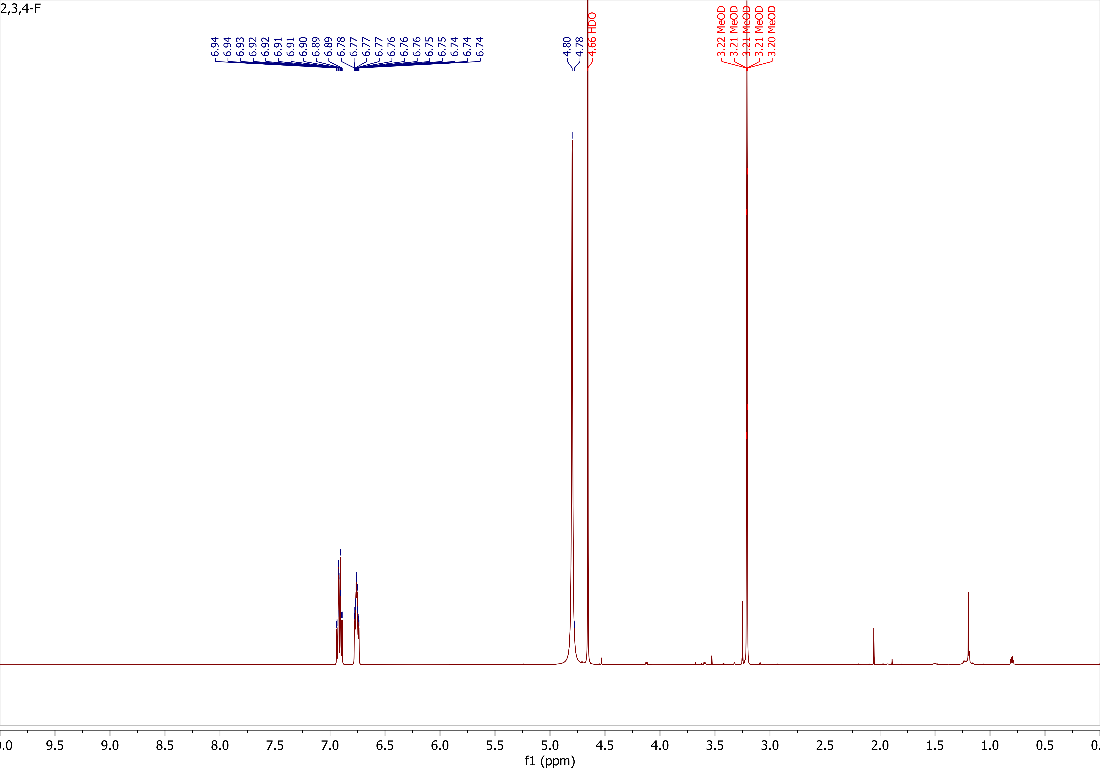

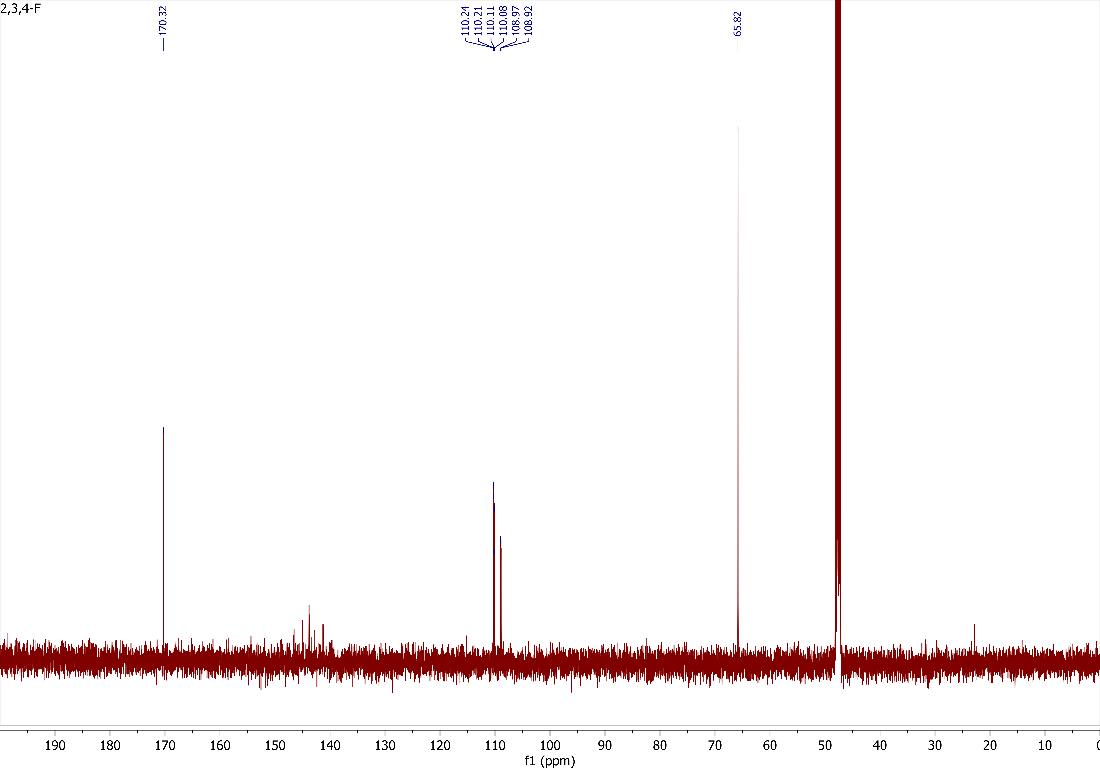

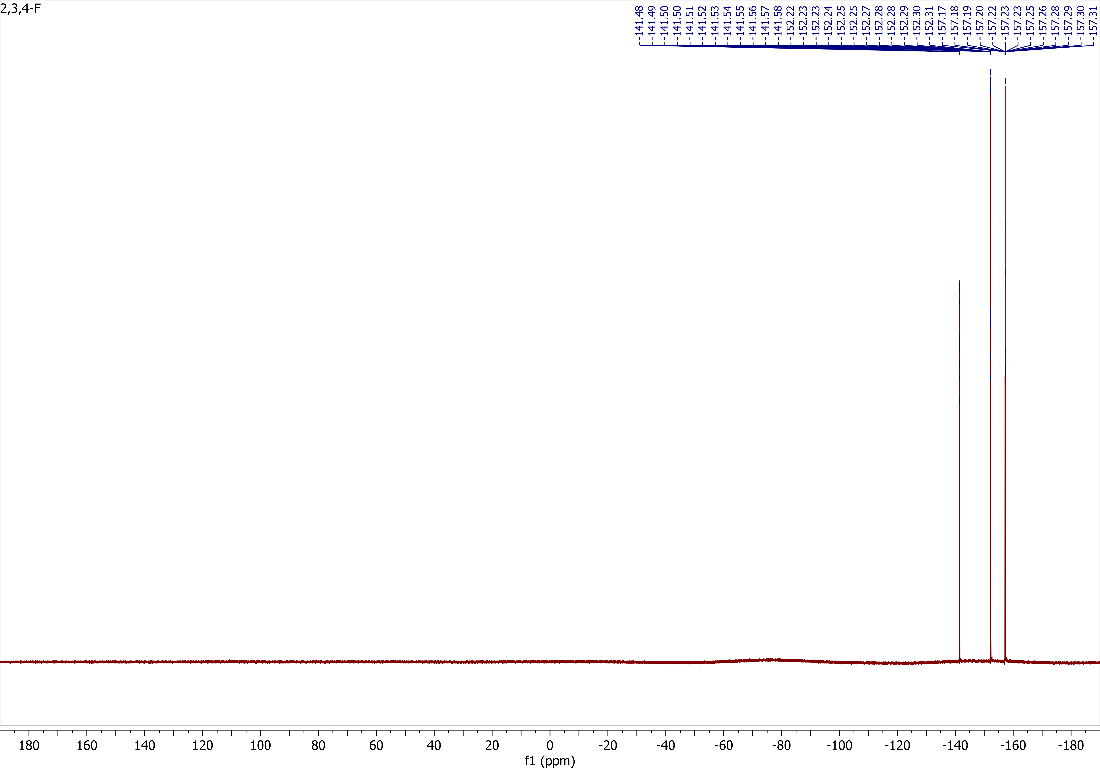

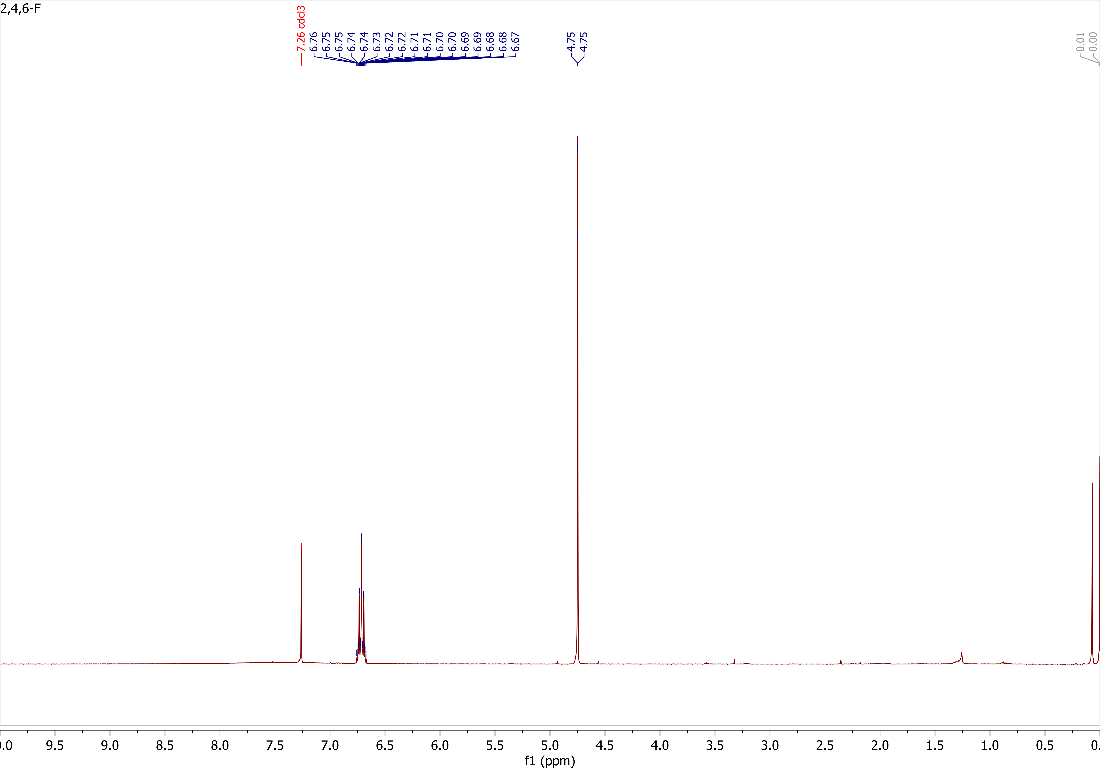

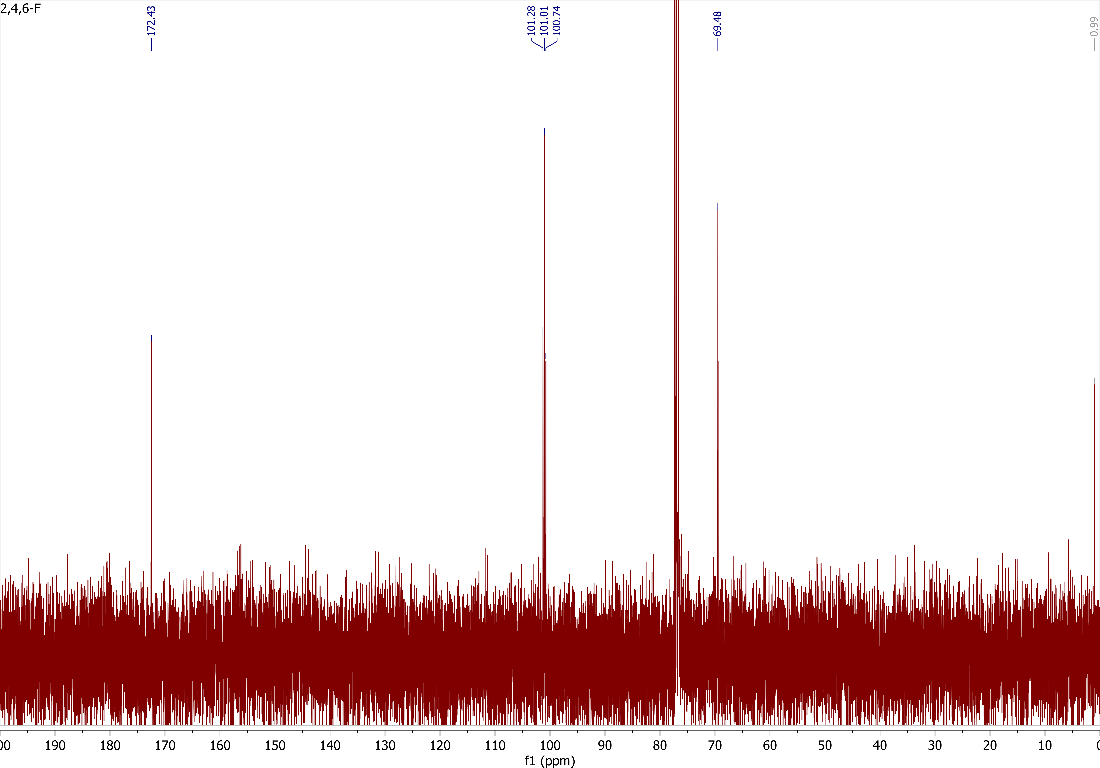

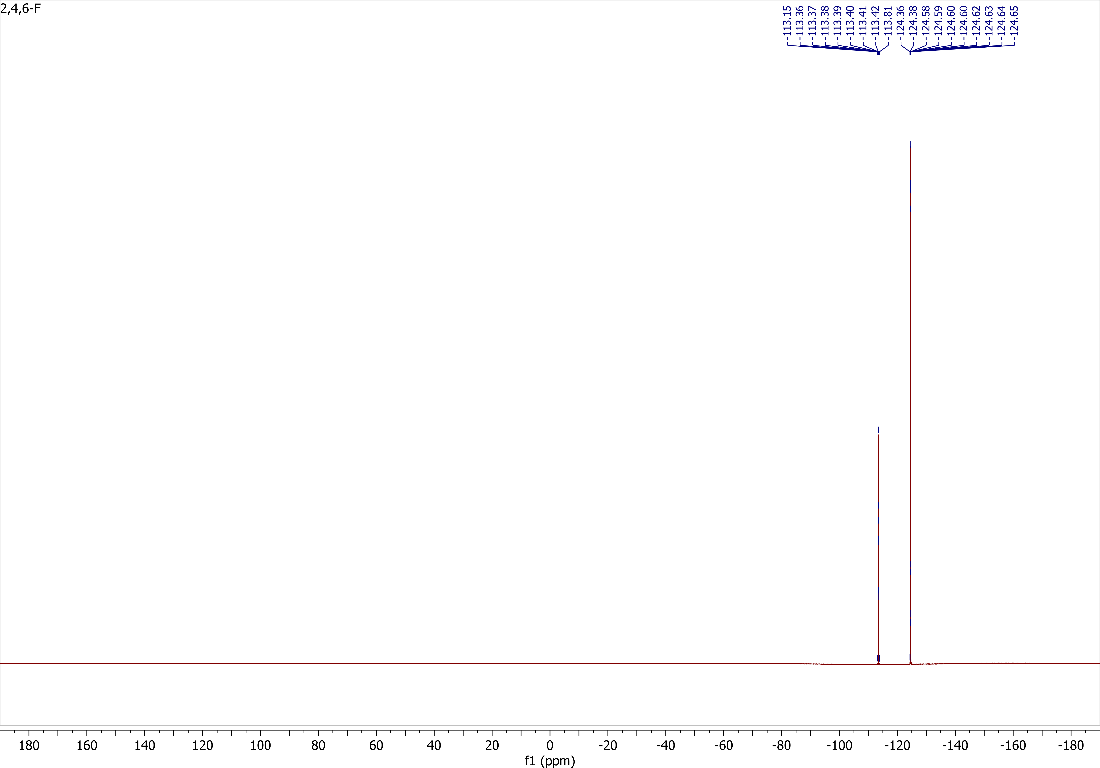

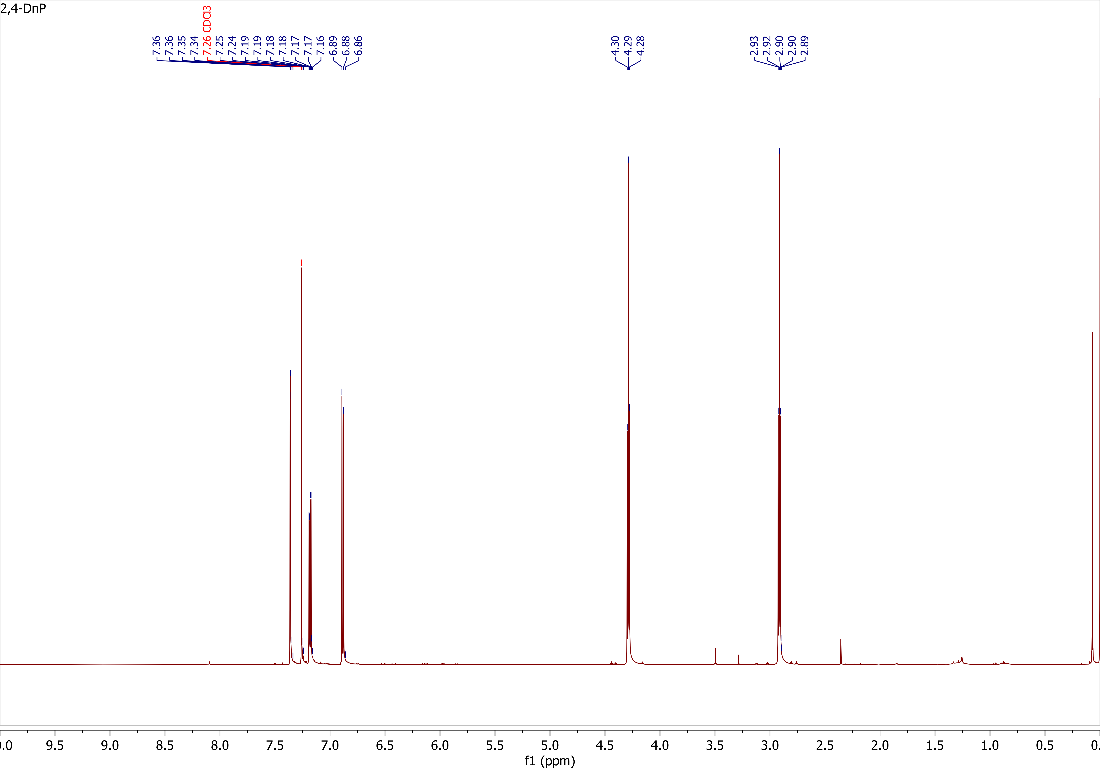

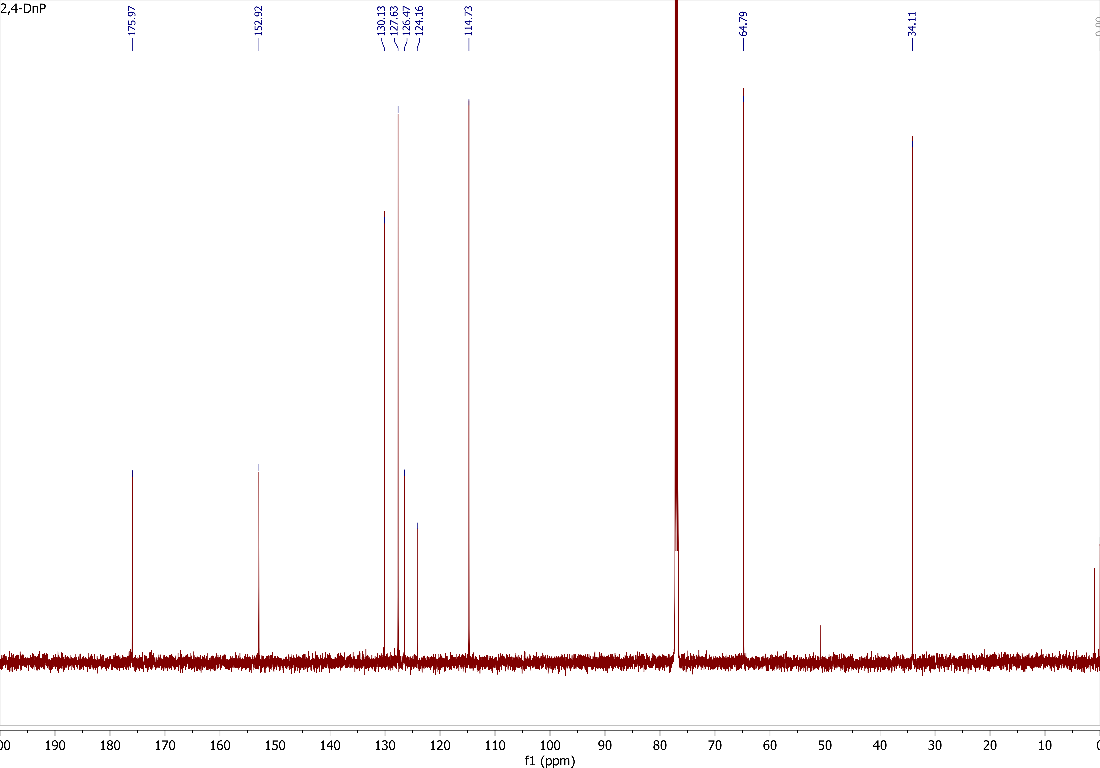
